# Scn4b Modulates Huntington’s Disease Phenotype Severity in vivo

**DOI:** 10.64898/2026.03.08.708251

**Authors:** Suphinya Sathitloetsakun, Vanessa Farrell, S. Sebastian Pineda, Hyeseung Lee, Jung Hoon Shin, Francisco J. Garcia, Raleigh M. Linville, Manolis Kellis, Veronica A. Alvarez, Myriam Heiman

## Abstract

Although it has been known for over 30 years that CAG trinucleotide repeat expansions in the *HTT* gene are the cause of Huntington’s disease (HD), it is still not understood how these mutations lead to the loss of striatal spiny projection neurons (SPNs) and other vulnerable neuronal cell types in HD. Here we show that *SCN4B*, a gene that is enriched in neurons that influence motor function, including striatal SPNs, modulates HD-associated phenotypes *in vivo*. Loss of *Scn4b* in wild-type mice mimics and *Scn4b* overexpression in an HD mouse model rescues several HD-associated phenotypes, including motor and cognitive deficits. Single nucleus RNA sequencing (snRNA-seq) analysis reveals that loss of *Scn4b* replicates several HD-associated gene expression signatures in the striatum. Conversely, overexpression of *Scn4b* rescues HD gene expression signatures and improves various SPN electrophysiological properties in HD model mice. Taken together, our results implicate loss of *Scn4b* expression as an important contributor to HD pathogenesis and therapeutic target in HD.

## INTRODUCTION

Huntington’s disease (HD) is a fatal neurodegenerative disease characterized by progressive motor impairments, cognitive decline, and psychiatric symptoms, as well as a marked loss of spiny projection neurons (SPNs) in the caudate nucleus and putamen (striatum).^1,2^ HD is caused by CAG trinucleotide repeat expansions in exon 1 of the huntingtin (*HTT*) gene, which lead to an extended polyglutamine tract in the encoded mutant huntingtin protein (mHTT).^3^ Despite the known causal mutation in *HTT*, the molecular mechanisms underlying HD pathogenesis and neuronal cell death are not fully understood.

A two-step model of HD pathogenesis proposes that HD progression involves: somatic expansion of the inherited expanded CAG repeat allele beyond a critical pathogenic threshold (or thresholds), followed by downstream toxicity processes that lead to disease manifestation, processes that are potentially cell type and context specific.^4^ Recent HD genome-wide association studies (GWAS) have provided insights into many HD age-of-onset modifiers, including DNA repair genes in the mismatch repair (MMR) pathway that are known to modify the rate of somatic CAG repeat instability.^4^ Loss of function of several of these genes can halt the expansion of CAG repeats and block the emergence of phenotypes in HD mouse models.^5^ Furthermore, the expression of some of these genes, such as *MSH2* and *MSH3*, is higher in SPNs compared to other cell types in the striatum, suggesting that differential expression of MMR genes may contribute to heightened cell type vulnerability in this brain region.^6^

While MMR genes have been implicated in the first step in HD pathogenesis, less is understood about cellular processes underlying mHTT toxicity (i.e. the second step of pathogenesis). Gain-of-function toxicities, at either the DNA, RNA, or protein level, are likely contributors.^4,7^ Some examples include loss of proteostasis, mitochondrial dysfunction, altered synaptic transmission, and transcriptional dysregulation.^8^ However, which of these are drivers versus secondary contributors to mHTT toxicity is not yet fully clear. Recent HD GWAS have identified five additional loci, aside from those linked to MMR genes, that are linked to progression of clinical phenotypes, including *MED15*, *RRM2B*, *CCDC82*, *TCERG1*, and a second chr22 locus.^9^ None of these genes has a clear role in DNA repair, and thus they are hypothesized to either indirectly contribute to CAG expansion or to modify downstream mHTT toxicity processes.^7,10^

Additional insights into mediators of mHTT toxicity may be gained by considering other diseases that have shared HD-like striatal pathology, but in developmental ages - including diseases linked to infantile basal ganglia degeneration. In such cases, alterations to genes and pathways essential for SPN survival would result in pediatric manifestations of an HD-like pathology. Gene mutations that are causal forms of these diseases (e.g. Leigh Syndrome) are involved in mitochondrial function, immune signaling, and ion channel activities,^11^ cellular pathways that are also dysregulated in HD.^7,12^

Previous transcriptomic studies using HD knock-in mouse models with increasing CAG repeats have revealed several genes that are dysregulated in a CAG repeat-length-dependent manner, and which are also dysregulated in human HD samples.^12,13^ Of these genes, *SCN4B*, sodium channel beta subunit 4, is of particular interest given that it is one of the most significantly downregulated striatal genes across various HD studies^12–14^ and mutations in its binding partner, *SCN2A*, are causal for forms of infantile basal ganglia degeneration.^11^ *Scn4b* is highly expressed in striatal SPNs and several other HD-affected neuronal cell types that impact motor function, including Purkinje neurons and deep layer neurons of the motor cortex. SCN4B is a single transmembrane-spanning protein with an extracellular N-terminus containing an immunoglobulin-like domain and an intracellular C-terminus.^15^ It interacts with the alpha subunit of the sodium voltage-gated channels to facilitate sodium influx needed to initiate an action potential.^15^ Additionally, it has also been shown to be important for gating the resurgent current, thus controlling firing frequency, and in regulating the induction of spike-timing-dependent long-term depression (tLTD) in neurons.^16,17^ Besides its electrophysical roles, SCN4B has been implicated in cell adhesion-related activities, spine density regulation, and axonal fasciculation in cell culture studies.^15,18,19^ It has also been shown to regulate the hypnotic effects of ethanol and other sedative drugs.^20^ Despite these diverse cellular roles of SCN4B, its contribution to mHTT toxicity mechanisms has not been directly tested.

To investigate the roles of *Scn4b* in HD pathogenic mechanisms, we performed *in vivo Scn4b* genetic studies to test if *Scn4b* causally contributes to HD-associated disease phenotypes. We show that post-developmental reduction of *Scn4b* in wild-type mice is sufficient to induce HD-associated behavioral and transcriptional phenotypes and that overexpression of *Scn4b* in the zQ175 HD mouse model can conversely rescue several HD-associated behavioral and transcriptional phenotypes. Together, these data demonstrate that *Scn4b* bidirectionally modulates HD-associated phenotypes *in viv*o and may act as an early driver of HD toxicity.

## RESULTS

### Scn4b is one of the most CAG-length-dependent dysregulated genes in an HD context

To uncover SPN cell type-specific genes dysregulated in a CAG-length-dependent manner, we performed a linear regression analysis of genes dysregulated in both direct (dSPN) and indirect (iSPN) pathway SPNs.^12^ *Scn4b* was one of the most dysregulated genes in a CAG-dependent manner in both cell types in mice after *Pde10a*, a known pathological driver of HD disease phenotypes^12^ (**Figure 1A**). Given this, we sought to next determine the extent of *SCN4B* downregulation in human HD samples of different Vonsattel grades (grades 1-3) compared to pathologically normal (PN) control samples. Consistent with findings in HD model mice that *Scn4b* downregulation starts at an early disease stage,^18^ *SCN4B* was significantly downregulated in the caudate nucleus of human HD grade 1 samples, and the level of *SCN4B* reduction correlated with HD disease progression (**Figure 1B-C**). Given its striatal enrichment, CAG-length-dependent downregulation in HD mouse model tissue, and reduction in grades 1-3 human HD tissue, we hypothesized that *SCN4B* downregulation is an early and causal contributor in HD disease mechanisms.

**Figure 1.**
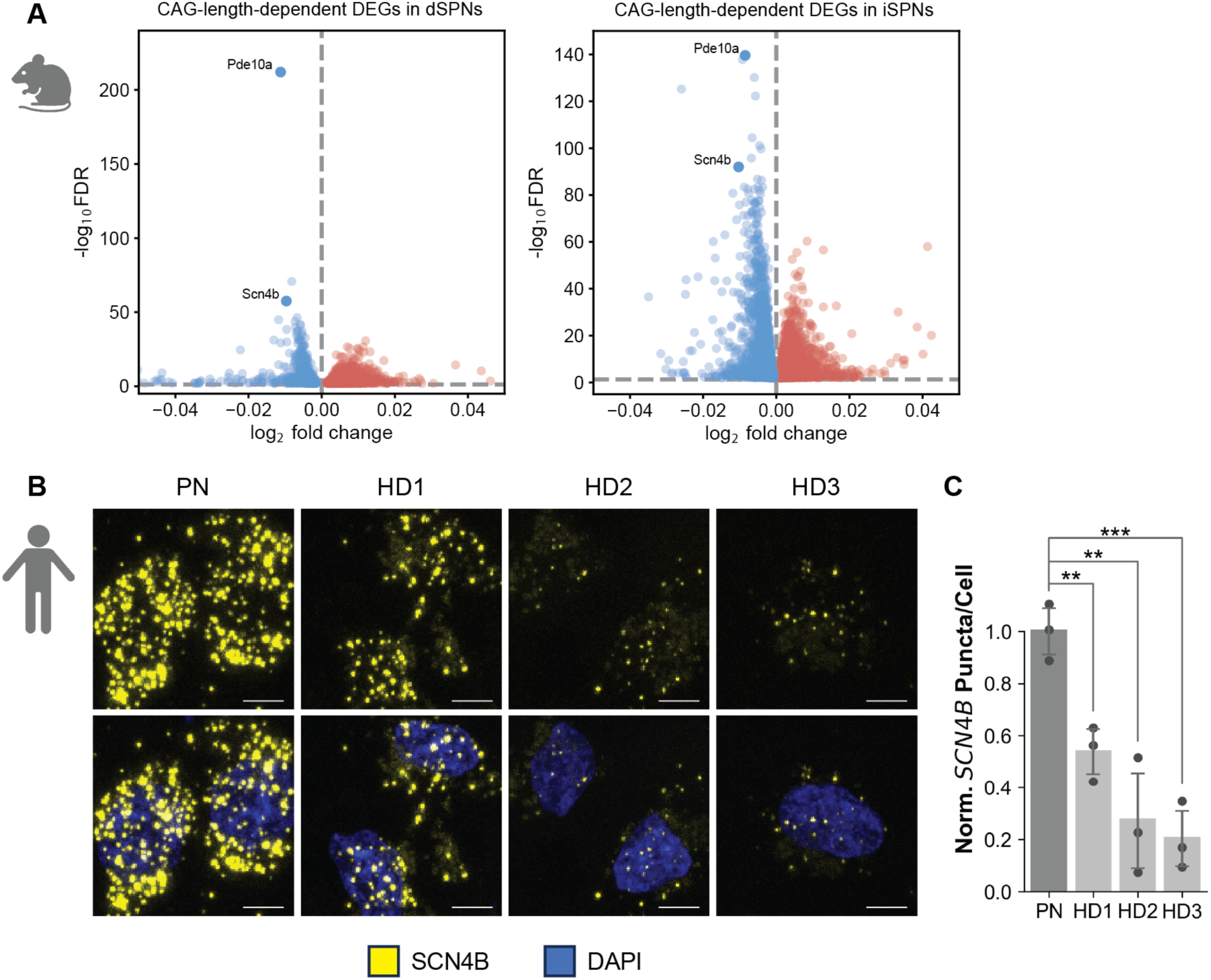
*Scn4b* is highly downregulated in HD mouse models and human HD SPNs. **(A)** Volcano plots of significantly differentially expressed genes (DEGs) in a CAG-length-dependent manner in the direct (dSPNs) and indirect (iSPNs) pathway spiny projection neurons of an HD mouse model allelic series (Q50, Q111, Q170, zQ175), as reported by translating ribosome affinity purification (TRAP) profiling.^12^ **(B)** Representative RNAScope *in situ* hybridization images showing striatal caudate nucleus *SCN4B* RNA (yellow) reduction across human HD grades 1-3 compared to pathologically normal (PN) caudate tissues. Scale bar, 5 µm. **(C)** Quantification of normalized (to PN) *SCN4B* puncta per cell. Data are shown as mean ± SD (*n*=3 biological replicates per condition; 10 images (63x) per donor). One-sided t-test: \*\**p* = 0.0031 (PN vs. HD1) and 0.0081 (PN vs. HD2); \*\*\**p* = 0.0007 (PN vs. HD3).

### Scn4b lowering in adult wild-type mice recapitulates several HD-associated phenotypes

To test the causal role of post-developmental *Scn4b* downregulation in contributing to HD-associated phenotypes, we performed an RNA knockdown (KD) study in adult wild-type (WT) mice, absent any mHTT context, using the CRISPR/CasRx system^21^ delivered via a blood-brain barrier-permeable AAV9 variant, PHP.eB^22^ (**Figure 2A**). We adopted this post-developmental approach for *Scn4b* KD (as opposed to germline knockout) to better mimic the disease associated *SCN4B* downregulation that occurs in HD and to overcome any known developmental compensation of *Scn4b* loss in mice.^19^ In this system, we designed an all-in-one AAV construct encoding CasRx with a 3xHA tag (CasRx-3xHA) under the control of the pan-neuronal human synapsin (hSyn) promoter, the woodchuck hepatitis virus post-transcriptional regulatory element 3 (WPRE3)^23^ to enhance expression, and three U6-driven guide RNAs (gRNA) targeting *Scn4b*. Because nuclear targeting of CasRx is required for efficient KD,^21^ we optimized the nuclear localization signal (NLS) for neurons, specifically striatal neurons. A combination of a 5′ SV40 NLS and a 3′ bipartite c-Myc NLS was necessary to achieve nuclear localization in striatal neuronal-like STHdh cells *in vitro* (**Figure S1A**), and was thus used in the final construct aiming to achieve nuclear localization in SPNs *in vivo*.

**Figure 2.**
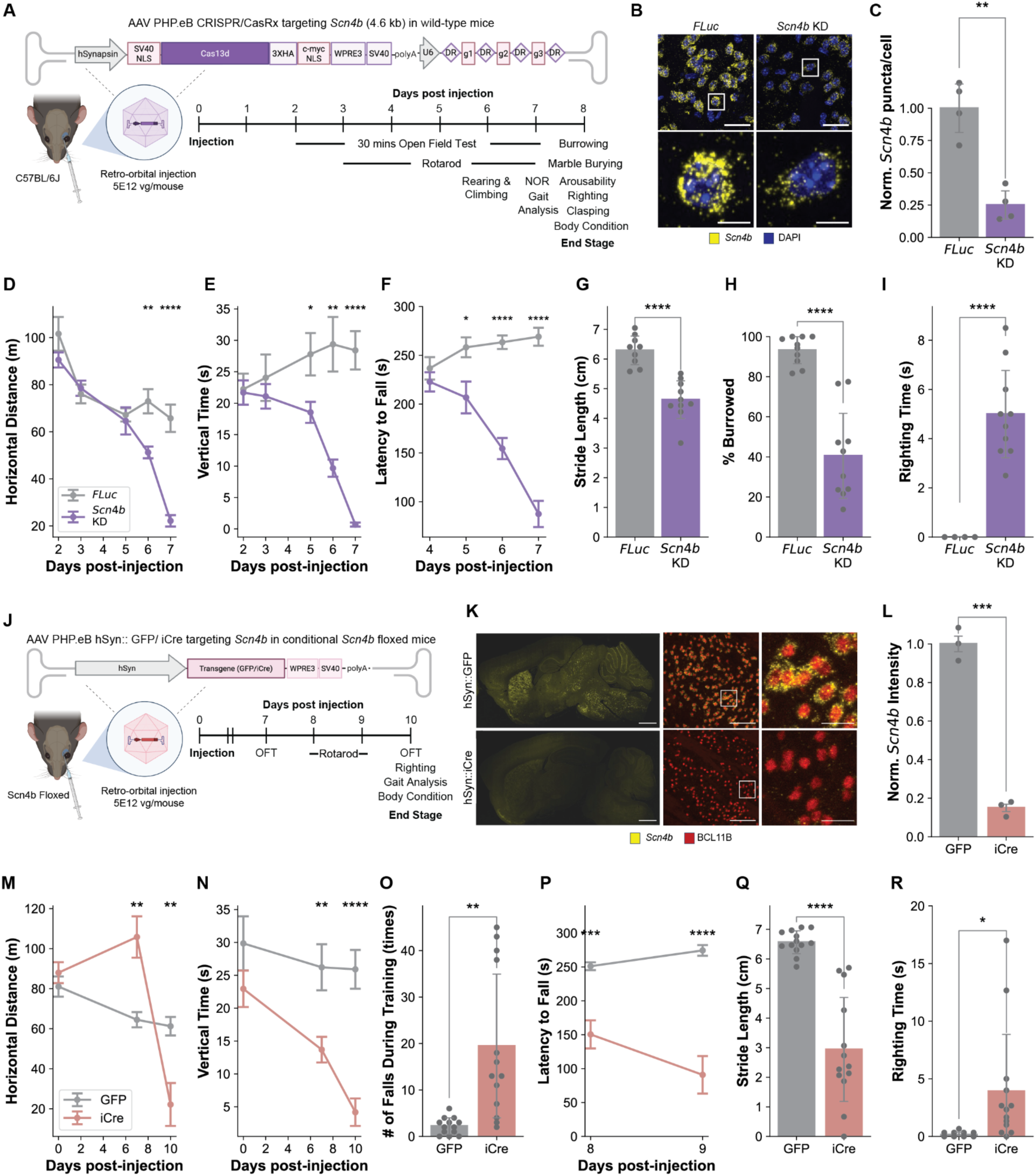
Post-developmental downregulation of *Scn4b* by either of two genetic *approaches* promotes HD-related motor and coordination deficits. **(A)** Schematic showing the *Scn4b* knockdown (KD) approach in adult wild-type mice and the experimental timeline for behavioral testing. **(B)** Representative RNAScope *in situ* hybridization images of striatal tissue from mice injected with 5E12 vg/mouse of AAV PHP.eB hSyn::CRISPR/CasRx containing guides targeting *Scn4b* showing the reduction of *Scn4b* signal (yellow). *FLuc* denotes data from tissue with the guides targeting Firefly Luciferase, which is not present in the mouse genome. Scale bar, 25 µm (top) and 5 µm (bottom). **(C)** Relative quantification of *Scn4b* puncta count. Data are shown as mean ± SD (*n*=4 mice per condition; 25 images (63x) per mouse). One-sided t-test: *p* = 0.0038. **(D)** Horizontal distance and **(E)** vertical time traveled as measured by open field test over a 30 min testing period. **(F)** Latency to fall in the rotarod test. **(G)** Stride length measured from paw prints from gait analysis. **(H)** Percentage of food pellets burrowed (by total mass) in the burrowing test. **(I)** Righting reflex time. Data are shown as mean ± SEM for all longitudinal time course data and mean ± SD for single-time-point comparisons (*n*=8-10 mice per group). One-sided t-test: \**p* ≤ 0.05, \*\**p* ≤ 0.01, \*\*\**p* ≤ 0.001, \*\*\*\**p* ≤ 0.0001. **(J)** Schematic showing the *Scn4b* knockout (KO) approach in conditional *Scn4b* floxed mice and the experimental timeline for behavioral testing. **(K)** Representative RNAScope *in situ* hybridization FISH images of striatal tissue from mice injected with 5E12 vg/mouse of AAV PHP.eB hSyn::GFP/iCre virus showing the reduction of *Scn4b* signal (yellow). Scale bar, 1 mm (left), 75 µm (middle) and 15 µm (right). **(L)** Relative quantification of *Scn4b* signal intensity. Data are shown as mean ± SD (*n* = 4 mice per condition; 10 images (40x) per mouse). **(M)** Horizontal distance and **(N)** vertical time traveled as measured by open field test over a 60 min testing period. **(O)** Number of falls during rotarod training and **(P)** latency to fall in the rotarod test. **(Q)** Measured stride length of the paw prints from gait analysis. **(R)** Righting reflex time. Data are shown as mean ± SEM for all line graphs and as mean ± SD for all bar graphs (*n*=13 mice per group). One-sided t-test: \**p* ≤ 0.05, \*\**p* ≤ 0.01, \*\*\**p* ≤ 0.001, \*\*\*\**p* ≤ 0.0001. Figures in panels A and J were created with BioRender.

Next, we designed gRNAs targeting *Scn4b*, assessed their KD efficacy *in vitro* using HEK293T cells expressing mouse *Scn4b* cDNA (**Figure S1B**), and selected the top three guides (maximum KD of 90% at the protein level and 80% at the RNA level) compared to the Firefly Luciferase (*FLuc*) non-targeting control; **Figure S1C-E**) to clone into the final construct in a pre-gRNA configuration,^21^ where the gRNAs are flanked by two direct repeats, and subsequently used this for virus production. To apply the system *in vivo*, we retro-orbitally injected 5E12 vg/mouse of PHP.eB-hSyn::CRISPR/CasRx virus containing validated guides targeting either *Scn4b* or the *FLuc* non-targeting control into 8-week-old C57BL/6J mice (*n*=10 per group, **Figure 2A**). We observed widespread neuronal expression of CasRx-3xHA throughout the brain, with ∼70% neuronal transduction in the striatum (**Figure S1F-H**), and there was no significant increase in microglial or astroglial activation as assessed by IBA1 (**Figure S1I-K**) and GFAP immunofluorescence (IF) staining (**Figure S1L-N**). Inspection of hSyn::GFP reporter expression by PHP.eB revealed slightly higher expression in matrix versus striosome SPNs by this viral approach (**Figure S1O**). Across all transduced neurons (i.e. SPNs expressing CasRx-3xHA), *Scn4b* RNA levels were reduced by ∼75% in the *Scn4b* KD mice compared to *FLuc* control KD mice as assessed by RNAScope *in situ* hybridization (**Figure 2B-C**). This level of decreased expression is comparable to the level of *Scn4b* reduction reported by our previous TRAP profiling of HD mouse models (90% in R6/2 and 62% in zQ175 mice),^12^ and reduction in HD grade 2-3 caudate nucleus tissue (75% in grade 2 and 80% in grade 3; **Figure 1B-C**).

While mice with the *FLuc* control KD perturbation exhibited no HD-mimetic phenotype, *Scn4b* KD mice displayed significant weight loss (*p* < 0.0001 at 8 days post injection; **Figure S2A**) with poor body condition scores (**Figure S2B**), and exhibited motor deficits similar to those characteristic of HD mouse models beginning at five days post injection. Due to the fast progression and severity of the phenotype, we performed a high-density measurement of HD-relevant behavioral tests to assess the motor and cognitive performance of these mice (**Figure 2A**). Open field testing was performed daily starting at 2 days post injection; by 7 days post injection, *Scn4b* KD mice became akinetic, with significant decreases in motor activities both horizontally and vertically (*p* < 0.0001; **Figure 2D-E and S2C-I**). Thigmotaxis analysis of horizontal activity time at 7 days post injection showed that *Scn4b* KD mice spent significantly less time in the center of the open field arena compared to *FLuc* control (*p* = 0.0343; **Figure S2J**).

Consistent with the observed decline in the open field vertical activity measurements, *Scn4b* KD mice also showed a reduction in the number of rearing and climbing activities (*p* = 0.0069; **Figure S2K**). On the rotarod test, despite their comparable performance to controls at the beginning of the testing on this assay at 4 days post injection (**Figure S2L)**, on all subsequent days *Scn4b* KD mice displayed a significant decrease in rotarod performance (*p* < 0.0001 after 7 days; **Figure 2F**). Similar to HD mouse models,^24^ *Scn4b* KD mice displayed an abnormal walking gait with shorter stride and stance lengths and non-overlapping fore and hind steps (*p* < 0.0001, *p* < 0.0001, and *p* = 0.0442, respectively; **Figure 2G and S2M-O**). *Scn4b* KD mice also showed deficits in the novel object recognition test (*p* = 0.0316; **Figure S2P**), the burrowing test (*p* < 0.0001; **Figure 2H**), and the marble burying test (*p* = 0.0002; **Figure S2Q**). Reflexive and sensorimotor responses were also impaired in *Scn4b* KD mice as they had a longer latency to right themselves once placed on their sides (*p* < 0.0001; **Figure 2I**), and to be aroused after cage opening (*p* = 0.0099; **Figure S2R**) and had higher hind-limb clasping scores (*p* = 0.0027; **Figure S2S**). A small fraction of *Scn4b* KD mice showed sudden death and seizure-like symptoms at the later stage of the experiment (**Figure S2T**). Due to this sudden death and given the known association of *SCN4B* with long QT syndrome and sudden cardiac death in humans^25^ as well as the possible trace expression of the hSyn promoter in cardiac tissue,^26^ we evaluated the *Scn4b* levels in bulk heart tissue. qPCR analysis revealed no difference in *Scn4b* level in the hearts of mice with *Scn4b* KD compared to *FLuc* control (**Figure S2U**), suggesting that the sudden deaths observed were not due to off-target cardiac effects of expression under the hSyn promoter. Similar behavioral deficits with a delayed onset of 12 days post injection were observed using a lower viral injection titer of 1E12 vg/mouse (*n*=4 per group; **Figure S3**) that resulted in 40% striatal neuronal transduction (**Figure S1G-N**) with 55% *Scn4b* reduction in these transduced neurons (**Figure S3B-C**). Overall, our results show that the post-developmental lowering of *Scn4b* alone via CasRx-mediated KD is sufficient to promote HD-related behavioral deficits.

To validate our findings using an orthogonal genetic approach, we performed a Cre-dependent *Scn4b* conditional knockout (KO) study, again absent the context of mHTT, in floxed mice that possess *LoxP* sites flanking exon 2 of *Scn4b*,^27^ which in the presence of Cre results in nonsense-mediated decay and loss of the *Scn4b* transcript (**Figure 2J**). We retro-orbitally injected 5E12 vg/mouse of AAV PHP.eB expressing either an improved Cre variant (iCre) or GFP control under hSyn promoter into 8-week-old *Scn4b* floxed mice (*n*=13 per group, **Figure 2J**). We confirmed using combined IF and RNAScope *in situ* hybridization that *Scn4b* was deleted in neurons across different brain regions in *Scn4b* KO mice, with 84% *Scn4b* reduction in the striatum 10 days post injection (**Figure 2K-L**). Strikingly, these *Scn4b* KO mice exhibited highly similar phenotypes within relatively the same timeline to those observed in the *Scn4b* CasRx KD mice, including significant weight loss (*p* = 0.0001; **Figure S4A**), poor body condition scores (**Figure S4B**), and severe motor deficits. In the open field test, *Scn4b* KO mice showed significant reductions in motor activity both horizontally and vertically at 10 days post injection (*p* = 0.018 horizontal distance, *p* < 0.0001 vertical time; **Figure 2M-N and S4C-H**). The mice also exhibited deficits in learning (*p* = 0.0004; **Figure 2O**) and performing the rotarod task (*p* < 0.0001; **Figure 2P**). *Scn4b* KO mice also displayed an abnormal walking gait with shorter stride and stance lengths and non-overlapping fore and hind steps (*p* < 0.0001, *p* < 0.0001, and *p* = 0.0440, respectively; **Figure 2Q and S4I-J)**. Additionally, *Scn4b* KO mice had a longer latency to right themselves when placed on their side (**Figure 2R**). Together, these results confirmed that brain-wide post-developmental *Scn4b* reduction (either via KD or conditional KO) is sufficient to promote HD-like behavioral deficits.

### Acute Scn4b knockdown mice share transcriptional signatures with HD models

To gain mechanistic insights into the basis of the observed acute *Scn4b* KD-associated phenotype toxicity, we performed single-nucleus RNA sequencing (snRNA-seq) on striatal tissue from the PHP.eB-hSyn::CRISPR/CasRx *Scn4b* KD and control mice, at the conclusion of behavioral testing, comparing *Scn4b* KD mice with *FLuc* non-targeting controls (*n*=6 mice per group; **Figure 3A**). We profiled 95,536 nuclei and annotated them into 22 transcriptionally distinct clusters including dSPN and iSPN striatal projection neurons, cholinergic and GABAergic interneurons, glial populations, progenitor cells, and vascular cell types based on the known markers (**Figure 3B**). SPNs were further characterized into matrix and striosome, and eccentric/outlier subclusters.^28^ UMAP visualization revealed good overlap between experimental groups and samples across major cell types (**Figure S5A-C**).

**Figure 3.**
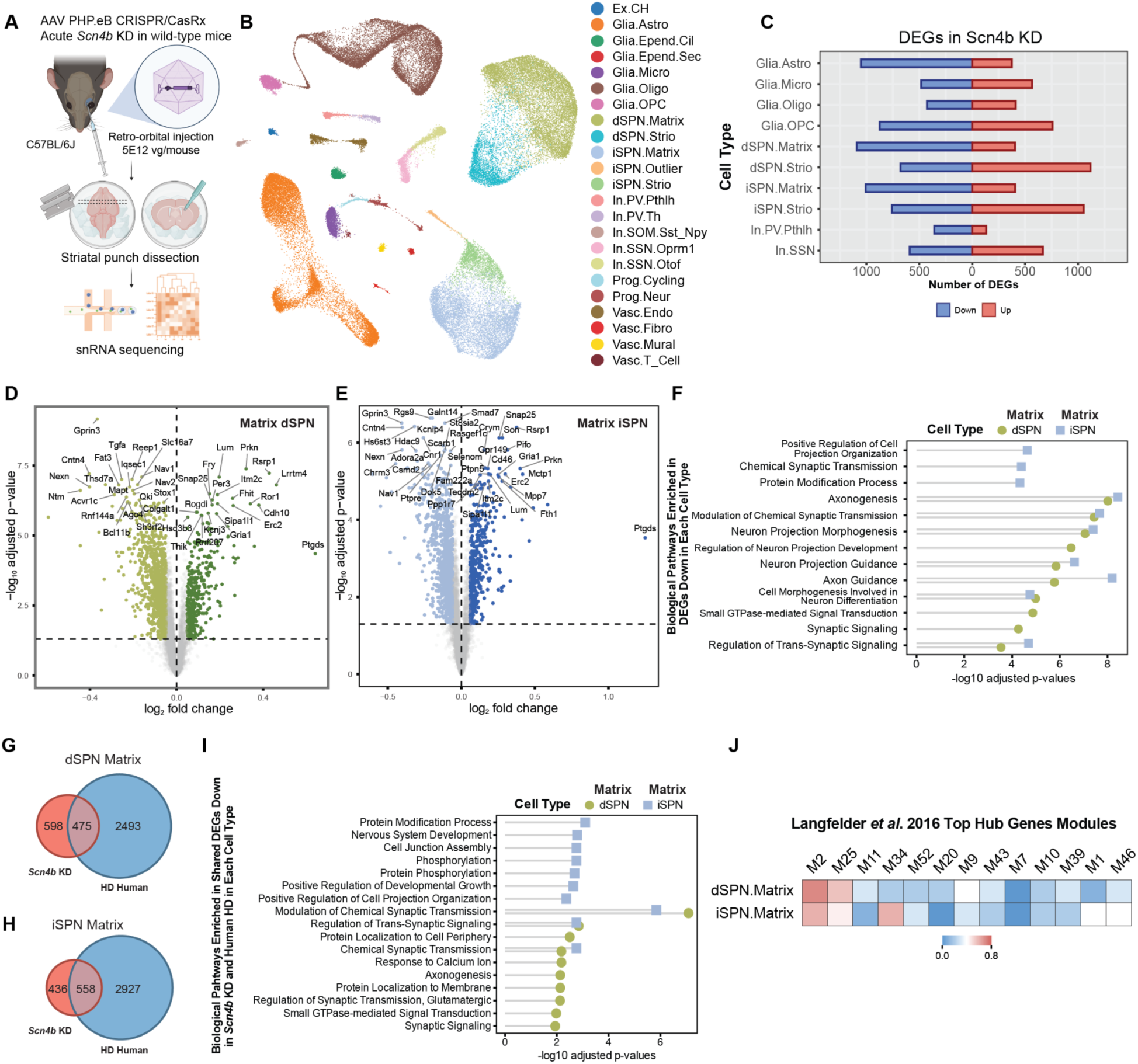
snRNA sequencing (snRNA-seq) analysis of striatal cell types reveals significant changes in dSPNs and iSPNs upon *Scn4b* KD. **(A)** Schematic showing the snRNA-seq experimental workflow. **(B)** UMAP representing all cell types captured from striatal tissue of the *Scn4b* KD and control (*FLuc*) mice. **(C)** Back-to-back bar chart showing the number of genes that are dysregulated in the striatum of *Scn4b* KD versus *FLuc* mice, per cell type (log_2_ fold change abs(*z*) >= 1 and *p* adj. FDR < 0.05). **(D)** Volcano plot showing significant DEGs (versus *FLuc*, log_2_ fold change abs(*z*) >= 1 and *p* adj. FDR < 0.05) in striatal matrix dSPNs and **(E)** iSPNs. **(F)** GO BP pathway enrichment analysis of *Scn4b* KD downregulated DEGs (versus *FLuc*) in matrix dSPNs and iSPNs. **(G)** Venn diagram showing the overlap between downregulated DEGs of *Scn4b* KD mice and human HD^12^ in matrix dSPNs and **(H)** iSPNs. Fisher’s exact test *p* = 1.32 × 10^-7^ and *p* = 1.61 × 10^-14^, respectively. **(I)** GO BP pathway enrichment analysis of the shared downregulated DEGs in matrix dSPNs and iSPNs from *Scn4b* KD mice and human HD. **(J)** Overlap of the *Scn4b* KD DEGs and top hub genes from Langfelder et al., 2016.^13^ Fisher’s exact test: *p* = 0.0097 (M2 dSPN.Matrix), 0.0771 (M2 iSPN.Matrix), 0.0337 (M25 dSPN.Matrix). Figure in panel A created with BioRender.

Pseudobulk DEG analyses identified many dysregulated protein-coding genes across various cell types upon *Scn4b* KD versus *FLuc* control. Striatal neuronal subtypes were among the cell types with the highest number of DEGs. Matrix SPNs showed more downregulated genes, with 1,091 downregulated and 405 upregulated genes in dSPNs, and 1,007 downregulated and 406 upregulated genes in iSPNs.

Striosome SPNs showed more upregulated genes, with 1,117 upregulated and 678 downregulated in dSPNs, and 1.053 upregulated and 758 downregulated in iSPNs [log_2_ fold change abs(*z*) >= 1 and *p* adj. FDR < 0.05 **(Figure 3C)**]. Given the higher KD viral expression in matrix compared to striosome SPNs by our gene knockdown approach **(Figure S1O)**, we focused our analysis on matrix SPNs. However, astrocytes and oligodendrocyte progenitor cells (OPCs) also showed many DEGs. In astrocytes, downregulated pathways were enriched in terms related to GTPase-mediated and intracellular signaling pathways, whereas upregulated pathways were enriched in ribosome biogenesis and oxidative phosphorylation. In OPCs, downregulated pathways were enriched in myelination and oligodendrocyte differentiation, whereas the upregulated pathways were enriched in synaptic and mRNA splicing processes. Given that hSyn does not drive expression in astrocytes or OPCs, these effects are likely secondary to neuronal changes upon *Scn4b* KD.

Of the DEGs in matrix SPNs, 638 downregulated genes and 234 upregulated genes were shared among the two cell types **(Figure S5D)**, indicating extensive shared DEGs across SPNs upon *Scn4b* KD. At the gene level **(Figure 3D-E)**, *Scn4b* KD resulted in downregulation of many components of the sodium channel complex that are known interacting partners of *Scn4b,* including the pore-forming α subunits *Scn8a* (Nav1.6) and *Scn3a* (Nav1.3), as well as another regulator of sodium channel function, *Fgf12*. Many ionotropic channel subunit encoding genes were also downregulated upon *Scn4b* KD, including those of potassium (*Kcna4, Kcnb2, Kcnd2, Kcnh4, Kcnip4*) and calcium (*Cacna1a/i, Cacna2d1/3*) channels. Beyond ion channel encoding gene expression changes, *Scn4b* KD also led to downregulation of various synaptic genes *(Homer1*, *Shank3*, *Nrxn3*, *Dlgap2/3*, *Cntn1/4/5*, and *Grip1*), as well as calcium-dependent activity signaling molecules (*Camk2b, Camkmt*). Gene ontology and pathway analyses also revealed shared downregulated DEGs in dSPNs and iSPNs of *Scn4b* KD mice to be enriched in terms related to synaptic signaling (modulation of chemical synaptic transmission, regulation of trans-synaptic signaling, IGSF CAM signaling), neuron projection processes (neuron projection morphogenesis, neuron projection guidance, cell morphogenesis involved in neuron differentiation), axon-related processes (axonogenesis, axon guidance), adherens junction, platelet activation and morphine addiction (**Figure 3F and S5E**), consistent with roles of Scn4b in modulating not only resurgent currents and action potential threshold, and cell adhesion and neurite outgrowth. Together, these findings support the known role of Scn4b in affecting intrinsic excitability and synaptic function in SPNs. In addition to these synaptic and cell adhesion genes, *Scn4b* KD also resulted in downregulation, across both SPN types, of many canonical SPN marker genes including *Pde10a*, *Rgs9*, *Cnr1,* and *Dclk3* as well as transcription factors enriched in and/or associated with SPN identity and function including *Foxp1*, *Bcl11b*, *Rarb*, *Rxrg*, and *Mn1*, genes that have been previously shown to be downregulated in HD and HD model tissue (**Figure 3D-E**).^12,13^

Shared upregulated DEGs in dSPNs and iSPNs of *Scn4b* KD mice included ionotropic and metabotropic channel-related genes, including *Gria1/2, Gabbr1*, *Kcnip3*, and *Kctd1*, likely representing compensatory mechanisms in the context of *Scn4b* KD. *Scn4b* KD also caused the upregulation of genes required for protein translation and oxidative phosphorylation, likely reflecting altered SPN metabolic demand in the context of synaptic changes (**Figure 3D-E and S5F-G**).

Given these specific changes to transcription factors and gene signatures with *Scn4b* KD in an otherwise WT context, *Scn4b* loss is likely a contributor to the transcriptional dysregulation observed in SPNs in HD and HD models. Indeed, we observed many known HD dysregulated genes when comparing DEGs in the *Scn4b* KD dataset to those of the HD STR266 genes dataset,^29^ including 47 and 48 downregulated DEGs (from 195 total downregulated genes) in dSPN and iSPN, respectively. To more fully assess the similarities between SPN dysregulated genes in *Scn4b* KD and HD mouse models, we compared the striatal DEGs of *Scn4b* KD mice to those previously reported for two HD mouse models: zQ175 and R6/2.^28^ Among the SPN-downregulated DEGs, we observed that 32.54% in dSPN or 32.08% in iSPN of *Scn4b* KD DEGs overlapped with DEGs in either the R6/2 or zQ175 HD model DEGs, or both (**Figure S5H-I**), a high amount given that these datasets each only represent one model timepoint. Gene ontology and pathway analyses revealed these shared DEGs to be enriched in pathways related to synaptic signaling (modulation of chemical synaptic transmission, regulation of synaptic transmission glutamatergic, chemical synaptic transmission, regulation of signal transduction, glutamatergic synapse, CGMP-PKG signaling pathway), and morphine addiction (**Figure S5J-K**). Lastly, we compared *Scn4b* KD DEGs to those previously reported for HD human SPNs. We found a 44.26% and 56.14% overlap between downregulated DEGs of *Scn4b* KD and human HD SPNs^28^ in dSPN and iSPN, respectively (**Figure 3G**). At the pathway level, these genes were highly enriched in synaptic signaling (modulation of chemical synaptic transmission, regulation of trans-synaptic signaling, chemical synaptic transmission, oxytocin signaling pathway, long-term depression), axon guidance, adherens junction, and morphine addiction (**Figure 3I and S5L**). Moreover, we also found great overlap within striatal-specific HD-relevant M2 and M25 modules when comparing to prior WGCNA analysis of HD model mice (**Figure 3J**). Taken together, these results demonstrate that *Scn4b* KD alone in an otherwise WT context recapitulates several key transcriptional signatures present in HD and HD mouse model striatal SPNs, including downregulation of synaptic genes, striatal marker genes, and genes associated with striatal transcriptional programs (e.g. *Foxp1*, *Bcl11b*, *Rarb*, *Rxrg*).

### Scn4b overexpression in zQ175 HD mice rescues HD model associated behavioral deficits

Given the causal role of *Scn4b* in HD mechanisms as implicated by our KD and KO studies, we next examined whether overexpressing (OX) *Scn4b* specifically in the striatum and motor cortex, both highly affected regions in HD, could ameliorate HD-related behavioral deficits in HD model mice (**Figure 4A**). For this purpose, we retro-orbitally injected 1E12 vg/mouse of AAV PHP.eB virus containing the full-length *Scn4b* coding sequence or GFP control under the control of a human *GPR88* MiniPromoter, an abbreviated version of the promoter that has been shown to give high expression in striatal projection neurons and pyramidal neurons in the motor cortex,^30^ into 2-month-old heterozygous zQ175 mice and their WT control littermates (*n*=6-7 per group). *Scn4b* OX increased the striatal *Scn4b* RNA level by 1.49 fold (from 0.55 to 0.82 relative to WT GFP; 49% increased relative to the disease level) in zQ175 *Scn4b* OX mice compared to the zQ175 GFP control (**Figure 4B-C**) after 11 months post injection (likely representing a lower overexpression than at peak expression as the AAV-mediated expression may decline over time).

**Figure 4.**
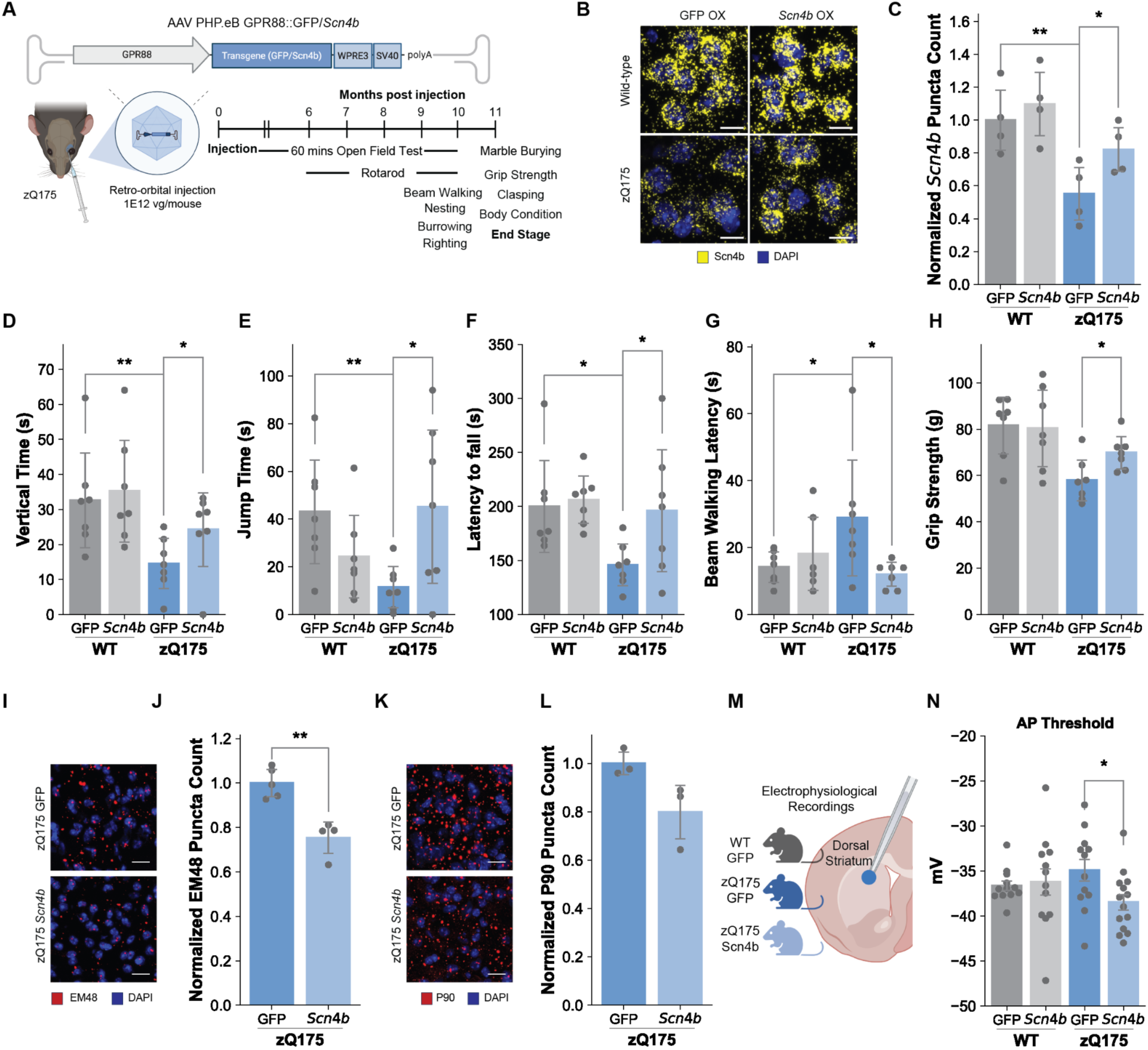
Striatal and motor-cortex-specific *Scn4b* overexpression in the zQ175 HD model mice rescues HD-model associated behavioral and cellular phenotypes. **(A)** Schematic showing the *Scn4b* OX approach in the zQ175 HD model mice and the timeline used for behavioral testing. **(B)** Representative RNAScope *in situ* hybridization images of striatal tissue from mice injected with 1E12 vg/mouse of AAV PHP.eB *GPR88*::GFP or *GPR88*::*Scn4b* OX showing the reduction of *Scn4b* (yellow) in zQ175 vs. WT-GFP OX and the overexpression of *Scn4b* in only zQ175-*Scn4b* vs GFP OX. Scale bar, 10 µm. **(C)** Relative quantification of *Scn4b* expression across the experimental groups. Data are shown as mean ± SD (*n*=3 mice per condition; 25 images (63x) per mouse). **(D)** Vertical time and **(E)** jump time parameters as measured by open field test over a 60 min testing period. **(F)** Latency to fall in the rotarod test. **(G)** Latency to cross in the beam walk test. **(H)** Grip strength measurement. For all tests, data are shown as mean ± SD (*n*=7 mice per group). One-sided t-test for all parameters: \**p* ≤ 0.05, \*\**p* ≤ 0.01. **(I)** Representative IF images (targeting EM48) of striatal sections from zQ175 mice injected with 1E12 vg/mouse of AAV PHP.eB *GPR88*::GFP or *GPR88*::*Scn4b*, showing the reduction of EM48+ mHTT aggregate signal (red) in *Scn4b* OX zQ175 mice. Scale bar, 20 µm. **(J)** Quantification of EM48+ mHTT aggregate count. Data are shown as mean ± SD (*n*=4 mice per condition; 15 images (40x) per mouse). **(K)** Representative IF images (using the P90 antibody) of striatal sections from zQ175 mice injected with 1E12 vg/mouse of AAV PHP.eB *GPR88*::GFP or *GPR88*::*Scn4b*, showing the reduction of P90+ mHTT aggregate signal (red) in *Scn4b* OX zQ175 mice. Scale bar, 20 µm. **(L)** Quantification of P90+ mHTT aggregate count. Data are shown as mean ± SD (*n*=3 mice per condition; 25 images (63x) per mouse). **(M)** Schematic representation of whole-cell clamp recording from the dorsal striatum. **(N)** The mean ± SEM of action potential thresholds for dorsal striatum SPNs from mice of the indicated genotypes. Figure in panel A created with BioRender.

While zQ175-GFP OX mice exhibited a significant decrease in motor activities and coordination as well as other behavioral deficits compared to WT-GFP OX control mice as expected at 13 months of age, these deficits were partially rescued with *Scn4b* OX in this model, without changes in body weight or body condition score between the zQ175 groups (**Figure S6A-B**). Specifically, in the open field test at 10 months post injection, zQ175-*Scn4b* OX mice showed a significant improvement in vertical and jump parameters (*p* = 0.0433 and *p* = 0.0144; **Figure 4D-E**) compared to the zQ175-GFP OX control mice. Other open field behavioral measures showed smaller significance or only non-significant improvement trends between groups (**Figure S6C-H**). They also exhibited significantly improved performance in the rotarod test (*p* = 0.016; **Figure 4F**) and beam walking assay (*p* = 0.0336; **Figure 4G**) compared to zQ175-GFP OX control mice. zQ175-*Scn4b* OX mice also had stronger grip strength (*p* = 0.0117; **Figure 4H**) and increased nesting activity (*p* = 0.0071; **Figure S6I-J**) compared to the zQ175-GFP OX controls. The marble burying assay, burrowing assay, righting test, and hindlimb clasping test were not significantly affected by *Scn4b* OX in zQ175 mice (**Figure S6K-O**). Taken together, these behavioral testing results demonstrated that long-term striatal and motor-cortex overexpression of *Scn4b* only is sufficient to rescue or improve several, but not all, HD-related motor deficits in the zQ175 HD mouse model.

### Scn4b overexpression decreases mutant huntingtin aggregates in zQ175 HD mice

To assess a cellular correlate of the improved behavioral performance of the zQ175 mice upon *Scn4b* OX, we sought to determine the level of *mHTT* protein aggregation in striatal samples of these mice, as *mHTT* aggregation is associated with disease model progression. For this purpose, we performed indirect immunofluorescence staining using the EM48 anti-huntingtin protein antibody, which recognizes a particular aggregated state of the polyQ tracts of mHTT protein. We observed a significant decrease in EM48 counts in the striatum of zQ175-*Scn4b* OX mice compared to zQ175-GFP OX control mice (**Figure 4I-J**). We observed a similar trend, though not significant (*p =* 0.053) using an antibody directed against the C-terminal proline 90 neoepitope of huntingtin exon 1 protein HTT1a (P90, which detects monomeric, oligomeric and aggregated mutant HTT1a) (**Figure 4K-L**).^31^

### Scn4b overexpression affects the action potential threshold in zQ175 HD mice

To assess whether striatal *Scn4b* OX in zQ175 mice affects SPN electrophysiological properties, whole-cell current clamp recordings were conducted from SPNs in the dorsal striatum (**Figure 4M**). As expected from prior reports,^32^ SPNs from zQ175-GFP OX mice showed reduced membrane capacitances (t(22) =2.95, *p* =0.008; **Figure S7A**) more depolarized (less negative) resting membrane potentials (t(23) =2.42, *p* =0.02; **Figure S7B**), and a trend to increased input resistances (**Figure S7C**) compared to WT-GFP OX SPNs. Current steps were delivered to elicit action potentials (APs) and the very first action potentials from each cell at rheobase were further analyzed (**Figure S7D**). The peak amplitude of the APs were significantly lower in zQ175-GFP OX mice than in WT-GFP OX mice (t(22) =4.01, *p* =0.0006; **Figure S7E**). Though not significant, the AP threshold was slightly increased in zQ175 GFP OX cells compared to WT GFP OX cells (**Figure 4N**). *Scn4b* OX had no effect in WT mice; however, the AP thresholds of zQ175-*Scn4b* OX mice were significantly changed compared to the zQ175-GFP OX controls (t(24) =2.348, p =0.0275; **Figure 4N**). *Scn4b* overexpression restored the AP threshold shift in zQ175 mice, supporting a causal model in which mHTT-driven loss of *Scn4b* moves the AP threshold away from the resting potential in SPNs. This finding is consistent with previous reports that *Scn4b* modulates AP thresholds in SPNs^16^ and that mHTT affects the excitability of SPNs.

### Scn4b overexpression reverses HD model transcriptional signatures

To investigate HD-associated transcriptional changes that might be rescued by *Scn4b* OX, we further performed snRNA-seq on striatal tissues from the four OX experimental groups: WT-GFP OX, WT-*Scn4b* OX, zQ175-GFP OX, and zQ175-*Scn4b* OX [*n*=6 mice per group; tissue harvested at 11 months post-injection (13 months of age)], after behavioral testing; **Figure 5A**). We profiled 300,157 nuclei and annotated them into transcriptionally distinct clusters including dSPN and iSPN striatal projection neurons, cholinergic and GABAergic interneurons, glial populations, progenitor cells, and vascular cell types (**Figure 5B)**. UMAP visualization revealed good overlap between experimental conditions, genotypes, and samples across major cell types (**Figure S8A-D**).

**Figure 5.**
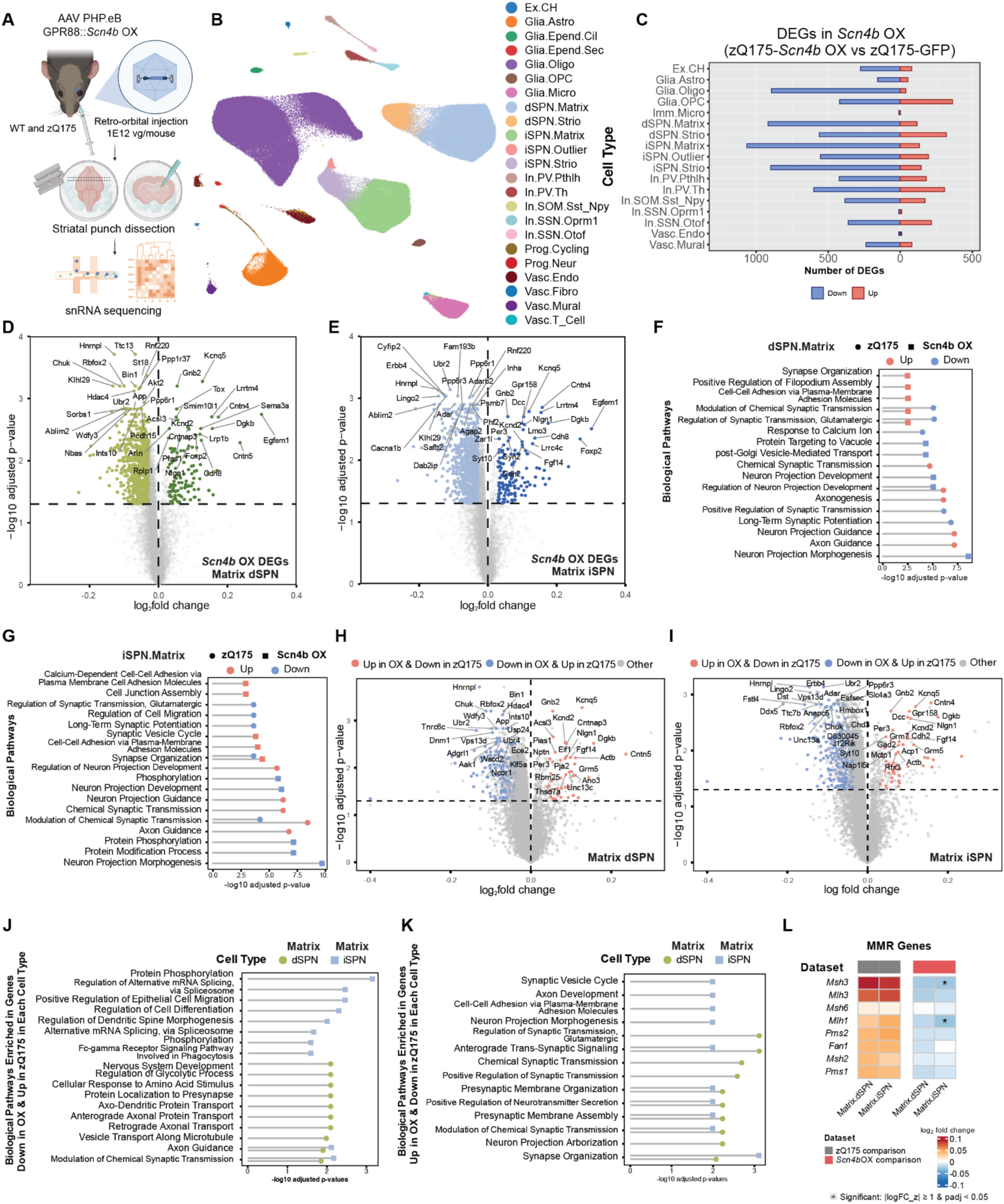
snRNA sequencing (snRNA-seq) analysis of striatal cell types in zQ175 mice upon *Scn4b* OX. **(A)** Schematic showing the snRNA-seq experimental workflow. **(B)** UMAP representing all cell types captured from striatal tissue of all mice included in this OX study. **(C)** Back-to-back bar chart showing the number genes that are differentially expressed (DEGs) in the striatum of *Scn4b* OX mice (zQ175-*Scn4b* OX vs zQ175-GFP OX; log_2_ fold change abs(*z*) >= 1 and *p* adj. FDR < 0.05), per cell type. **(D)** Volcano plot showing these genes in matrix dSPNs and **(E)** iSPNs of *Scn4b* OX mice. X-axis limit is set to ±0.4 for visualization purposes; genes exceeding this range include *Ttr* (log_2_ fold change = −0.778) and *Cmss1* (log_2_ fold change = −0.777) in dSPNs and *Cmss1* (log_2_ fold change = −0.818) in iSPNs. GO BP pathway enrichment analysis showing the upregulated and downregulated pathways of matrix **(F)** dSPNs and **(G)** iSPNs in the zQ175 and *Scn4b* OX comparisons. **(H)** Volcano plots showing genes that are changed in opposite directions in the zQ175 and Scn4b OX comparisons in matrix dSPNs and **(I)** iSPNs. GO BP pathway enrichment analysis of **(J)** genes that are downregulated in the *Scn4b* OX comparison and upregulated in the zQ175 comparison and **(K)** genes that are upregulated in the *Scn4b* OX comparison and downregulated in the zQ175 comparison. **(L)** Heatmap showing log_2_ fold change of selected MMR genes in the zQ175 and *Scn4b* OX comparisons. *indicates significance using the log_2_ fold change abs(*z*) >= 1 and *p* adj. FDR < 0.05 cutoff. Figure in panel A created with BioRender.

To determine the baseline transcriptional dysregulation due to the zQ175 model itself at 13 months in these mice, we first identified DEGs between zQ175 (zQ175-GFP OX) and wild-type control (WT-GFP OX) across various cell types (**Figure S8E**). We observed many DEGs (log_2_ fold change abs(*z*) >= 1 and *p* adj. FDR < 0.05) in both matrix dSPNs (538 downregulated and 783 upregulated genes) and iSPNs (495 downregulated and 824 upregulated genes) (referred to as zQ175 comparison hereafter; **Figure S8F-G**). These DEGs were consistent with previously published DEGs from zQ175 mice,^28^ confirming an expected HD-associated transcriptional signature in our dataset (**Figure S8H-I)**.

We next sought to assess the effect of *Scn4b* OX within the disease model context by comparing DEGs of zQ175 mice with *Scn4b* OX (zQ175-*Scn4b* OX) versus controls (zQ175-GFP OX) (**Figure 5C)**. We observed many DEGs in both matrix dSPNs (918 downregulated and 116 upregulated genes) and iSPNs (1,066 downregulated and 133 upregulated genes) (referred to as *Scn4b* OX comparison hereafter; **Figure 5D-E**). Among these, 592 downregulated and 66 upregulated were shared between the two SPN matrix cell types, suggesting a largely shared response in these cells (**Figure S9A**). The shared downregulated DEGs were enriched in pathways associated with synaptic signaling (modulation of chemical synaptic transmission, IGSF CAM signaling, phospolipase D signaling pathway), neuronal development processes (neuron projection morphogenesis, neuron projection development, plasma membrane bounded cell projection morphogenesis), vesicle trafficking (post-Golgi vesicle-mediated transport, endocytosis), and ATP-dependent chromatin remodeling (**Figure S9B-C**).

The shared upregulated genes were enriched in pathways associated with synaptic signaling/organization (modulation of chemical synaptic transmission, synapse organization), cell adhesion (cell-cell adhesion via plasma-membrane adhesion molecules, calcium-dependent cell-cell adhesion via plasma-membrane adhesion molecules) (**Figure S9D-E**). Several of these pathways were directionally opposite to those enriched in the zQ175 baseline comparison, suggesting that *Scn4b* OX affects several facets of mHTT-associated transcriptional dysregulation, in particular those related to synaptic signaling/organization (modulation of chemical synaptic transmission, regulation of synaptic transmission glutamatergic, IGSF CAM signaling, synapse organization) and regulation of neuron projection development (**Figure 5F-G, S9F-G**), as well as genes regulated by the Polycomb Repressive Complex (PRC), shown by prior studies to be dysregulated in an HD context^34–36^.

To identify rescued genes upon *Scn4b* OX in zQ175 mice, we focused on DEGs that show opposite directionality between the zQ175 (zQ175-GFP OX versus WT-GFP OX) and zQ175-*Scn4b* OX (zQ175-*Scn4b* OX versus zQ175-GFP OX) comparisons. The majority of these genes were upregulated in the zQ175 model but downregulated upon zQ175-*Scn4b* OX, with 187 genes in matrix dSPNs and 261 genes in matrix iSPNs (**Figure 5H-I)**. Among those, 133 genes were shared across the two cell types, including genes encoding subunits of sodium channels (*Scn8a*), potassium channels (*Kcnab2, Kcnc3, Kcnj9, Kcnn3, Kcns2*), calcium channels (*Cacna1b/h/i*), synaptic genes (*Nrxn2*, *Dlgap4*, *Syngap1*, *Unc13a*) as well as HD-implicated transcriptional regulators (*Tcerg1*, *Rbfox2*). At the pathway level, these genes are enriched in terms related to synaptic signaling (modulation of chemical synaptic transmission, IGSF CAM signaling), and axon guidance (**Figure 5J, S9H)**. A smaller set of rescued genes was downregulated in the zQ175 model but upregulated in the zQ175-*Scn4b* OX context, with 88 genes in dSPNs and 104 genes in iSPNs (**Figure 5H-I)**. Among those, 50 genes were shared across the two cell types, including channel-related genes and their regulators (*Fgf14*, *Kcnd2, Kcnq5*), synaptic related genes (*Nlgn1*, *Grik2*, *Grm5*, *Grm7*), calcium-dependent activity signaling molecule (*Camk2n1)*, and SPN enriched genes (*Arpp21*, *Cntn5*). At the pathway level, these genes are enriched in terms related to synaptic signaling (anterograde trans-synaptic signaling, modulation of chemical synaptic transmission, glutamatergic synapse, positive regulation of neurotransmitter secretion, neuroactive ligand signaling) and synaptic organization (synapse organization, presynaptic membrane organization, presynaptic membrane assembly) (**Figure 5K, S9I)**.

To investigate the relationship between *Scn4b* expression and regulators of somatic instability, we examined the expression of DNA MMR genes in our zQ175 baseline and *Scn4b* OX comparisons. A focused analysis of the MMR gene set revealed that several MMR genes displayed a trend to upregulation in the zQ175 dataset (consistent with our prior significant findings in the R6/2 mouse model^12^) and that they were in turn downregulated in the *Scn4b* OX dataset (**Figure 5L**). Overall, our data demonstrates that *Scn4b* OX alone in zQ175 SPNs partially restores the transcriptionopathy associated with mHTT, and notably not only with regard to synaptic gene expression.

## DISCUSSION

In this study we demonstrate that *Scn4b*, sodium channel beta subunit 4, contributes causally to HD-linked pathogenic mechanisms. Post-developmental *Scn4b* downregulation (by two independent genetic approaches, CRISPR/CasRx *Scn4b* KD and conditional floxed *Scn4b* mouse line KO) in adult WT mice (which mimics the downregulation seen in HD and HD models) was sufficient to recapitulate several HD-like behavioral deficits and transcriptional disease signatures in otherwise WT mice, while chronic *Scn4b* overexpression in the zQ175 HD model mice rescued several HD model-associated behavioral deficits, rescued SPN action potential (AP) shifts, reduced mHTT aggregates, and ameliorated mHTT-associated transcriptional dysregulation. Our findings on *Scn4b* are consistent with another study (Langfelder, Wang, et al. BioRxiv, 2026) showing that *Scn4b* reduction has profound effects on the mHTT-associated striatal transcriptome, mHtt aggregation, and patient-derived HD SPN cell death.

Mechanistically, Scn4b modulates sodium channel and thus neuronal electrophysiological properties, with prior work showing that its KD or KO in SPNs leads to a more depolarized AP thresholds, as well as a impaired spike-timing-dependent long-term depression and for backpropagating APs to evoke dendritic calcium transients.^16,17^ Altered SPN excitability and calcium transients would in turn affect various important downstream transcriptional regulators, especially those that are calcium regulated, and directly lead to transcriptional dysregulation in SPNs. Our data collectively show that these events downstream of *Scn4b* downregulation play a causal role in generating HD physiological phenotypes, and predict that a slow decrease to *Scn4b* expression (as opposed to acute KD/KO as performed here) may provide a useful new model of mHTT toxicity for therapeutic testing. Our data also show that *Scn4b* OX alone in HD model mice is sufficient to rescue many HD-associated changes.

*Scn4b* downregulation may thus be a critical component of the hypothesized second step in HD pathogenesis (cellular toxicity after CAG repeat expansion beyond a certain threshold), and this raises the possibility that effective gene therapies independent of those targeting DNA MMR and *mHTT* CAG somatic instability could target sodium channel activity and/or SPN electrophysiological abnormalities. Interestingly, *Scn4b* OX led to a decrease in MMR gene expression in zQ175 mice (**Figure 5**), and thus it is possible that such a therapeutic could also address the first step of HD pathogenesis.

Aside from having a casual role in HD, *SCN4B* may also have potential roles in the etiology of other neurodegenerative diseases. Specifically, high *SCN4B*-expressing cortical projection neurons of the cortex have recently been shown by our profiling to be one of the most affected cell populations in ALS and FTLD^37^, prompting future work to study SCN4B-dependent effects in the causal mechanisms underlying ALS/FTD. *SCN4B* might also contribute to mechanisms involved in the pathology of other neurological disorders, including Leigh syndrome^11^ and Autism Spectrum Disorder^38^, as these diseases are associated with mutations in *SCN2A*, of which SCN4B is an interacting partner. Future work aimed at understanding the transcriptional and functional regulation of *SCN4B* in HD, as well as these other brain-related diseases, will provide more insights into the molecular mechanisms underlying selective vulnerability and disease toxicity in various disease contexts and potentially offer new therapeutic targets.

## RESOURCE AVAILABILITY

### Lead contact

Further information and requests for resources and reagents should be directed to the Lead Contact, Myriam Heiman.

### Materials availability

This study did not generate new unique reagents.

## EXPERIMENTAL MODEL AND STUDY PARTICIPANT DETAILS

### Animals

All animal husbandry and experimental procedures were conducted with the approval of MIT and the NIH/National Institute on Alcohol Abuse Animal Care and Use Committees. Mice were group-housed under standard conditions with food and water provided *ad libitum* on a standard 12 hr light/dark cycle in pathogen-free facilities. No procedures were performed on the mice prior to the outlined experiments. Mice were obtained from the Jackson Laboratory (Bar Harbor, ME) or from breeding at MIT. Male C57BL/6J wild-type mice (Jackson Laboratory stock #000664) were used at 8-9 weeks of age for the CRISPR/CasRx KD studies. Male *Scn4b* floxed mice B6.129(SJL)-*Scn4btm1.1Geh*/RahJ (Jackson Laboratories stock #027530) were used at 8 weeks of age for the *Scn4b* conditional KO study. Female zQ175 HD model and their wild-type control littermates (B6J.zQ175DN KI, Jackson Laboratory stock #029928) were used at 8 weeks of age for the *Scn4b* OX study. For all studies, mice were randomly assigned to experimental groups. Behavioral experiments were performed at the time points outlined in the Method Details. In some instances, few mice were re-sorted after the first (baseline) open field test prior to AAV injection to ensure no difference in baseline behaviors among the experimental groups. Once *Scn4b* KD mice or zQ175 HD model mice became morbid, access to hydrated diet DietGel76A (ClearH2O, Portland, ME) was provided to all cages in the given experiment for additional nutritional support.

### Cell lines

HEK293T/17 cells were obtained from ATCC and striatal derived STHdh Q7/7 cells were obtained from the Coriell Institute. Cells were maintained according to the manufacturers’ recommended culture protocol. HEK293T/17 cells were cultured in DMEM with GlutaMAX, pyruvate, and non-essential amino acids (Gibco) with 5-10% fetal bovine serum (Gemini) at 37°C with 5% CO₂ and were passaged with 0.05% Trypsin-EDTA every 2-3 days at a ratio of 1:8. STHdh Q7/7 cells were cultured in DMEM with GlutaMAX and 10% fetal bovine serum (FBS) at 37°C with 5% CO₂ and were passaged once they reached 80% confluency at a ratio of 1:2 with the maximum passage number of P14.

### Human samples

Human tissue analyses were conducted as exempt human research, as this was secondary research using biological specimens not specifically collected for this study. All samples were obtained from biobanks/repositories as follows, using appropriate de-identification and under consent. Human post-mortem caudate nucleus tissue samples from pathologically normal (PN) controls and patients with HD were obtained from the Harvard Brain Tissue Resource Center, NIH Neurobiobank, Netherlands Brain Bank, and the University of Alabama at Birmingham. HD samples were examined by a neuropathologist and assigned pathological grades of 1, 2, 3, or 4.

## METHOD DETAILS

### Constructs cloning and guide design

The pAAV hSyn::CRISPR/CasRx construct was cloned using pAAV hSyn_Phos_CA_Citrine, a gift from Simon Wiegert (Addgene plasmid #98218), as a vector backbone. Using traditional restriction enzyme digestion and DNA ligation methods, sequences following the hSyn promoter up to the polyA signal were replaced with the CasRx - U6::guide sequences amplified from pAAV EF1-a-CasRx, a gift from Patrick Hsu (Addgene plasmid #109049) and WPRE3 sequences amplified from pAAV-CW3S-EGFP, a gift from Bong-Kiun Kaang (Addgene plasmid #61463). The 3xHA epitope tag was amplified from pAAV-CMV-SaCas9-3XHA, a gift from Feng Zhang (Addgene plasmid #61591) and later cloned into the construct using Gibson Assembly (New England Biolabs, Ipswich, MA). All linker sequences between these elements were removed using Q5 Site Directed Mutagenesis Kit (New England Biolabs, Ipswich, MA). To build the constructs for nuclear localization signal (NLS) testing, a 3’ SV40 NLS was replaced with a c-Myc NLS amplified from pLenti-c-Myc-Puro GFP, a gift from Fiovanni Tonon (Addgene plasmid #12329). Guides targeting *Scn4b* were designed using the CHOPCHOP web tool ^39^ with the option to target the gene in the *Mus musculus* (mm10/GRCm38) genome utilizing the CRISPR/Cas13 system for knockdown with a specific targeting sgRNA length of 30 nucleotides. Top guides based on the algorithm on/off target scoring were selected for each target and individually cloned into the pAAV hSyn::CasRx vector. Specifically, guide oligos were annealed using NEB T4 Ligation kit (New England Biolabs, Ipswich, MA) and cloned into the vector using BbsI sites preceding the U6 promoter. These constructs were used for *in vitro* validation experiments and the three best guides with maximum knockdown were then selected for each gene. To clone the final knockdown construct for each target, a guide array of the three top validated guides was synthesized with 5’ caagtaaacccctaccaactggtcggggtttgaaac 3’ spacer sequence linking each guide. The sequence of the *FLuc* non-targeting control array was as follows: 5’ CTACCTGGTAGCCCTTGTATTTGATCAGGCcaagtaaacccctaccaactggtcggggtttgaaacTGCCACTACTGTTCATGATCAGGGCGATGGcaagtaaacccctaccaactggtcggggtttgaaacTGCGCCAGGCTTGTCGTCCCCTTCGGGGGT 3’.^21^

The pAAV *hSyn*::iCre construct was cloned using pAAV.hSyn.eGFP.WPRE.bGH, a gift from James M. Wilson (Addgene plasmid #105539), as a vector backbone. A traditional restriction enzyme digestion and DNA ligation approach was also used to replace the GFP fragment with the iCre fragment. The iCre fragment was amplified from pENN.AAV.hSyn.Cre.WPRE.hGH, a gift from James M. Wilson (Addgene plasmid #105553). All constructs were sequence verified using whole-plasmid sequencing prior to the *in vitro* validation experiments and downstream virus production.

The pAAV *GPR88*::GFP construct was cloned using pAAV hSyn::GFP as a vector backbone. Similar to the above, a traditional restriction enzyme digestion and DNA ligation approach was used to replace the *hSyn* promoter fragment with the *GPR88* MiniPromoter. The *GPR88* MiniPromoter was amplified from pRMS1995, a gift from Elizabeth Simpson (Addgene plasmid #49125). The GFP sequence was replaced with *Scn4b* sequence, amplified from the mouse *Scn4b* cDNA ORF plasmid (Scn4b #MR226154; Origene, Rockville, MD) for the pAAV *GPR88*::*Scn4b* OX construct.

### Transient transfection for CasRx nuclear localization and guide RNA validation

For nuclear localization testing, HEK293T (ATCC, Manassas, VA) and STHdHQ7/7 cells (Coriell Institute, Camden, NJ) were seeded onto poly-D lysine-coated coverslips (VWR, Radnor, PA) using Dulbecco Modified Eagle’s Medium (DMEM) containing L-glutamax (ThermoFisher Scientific, Rockford, IL) supplemented with 10% FBS (Gemini, West Sacramento, CA) in 24-well plates 24 hr prior to transfection. Cells were transfected with 1 μg of the AAV CasRx constructs containing a 3’ SV40 NLS or a 3’ c-Myc NLS flanking the CasRx sequence using Lipofectamine3000 reagents (ThermoFisher Scientific, Rockford, IL) according to the manufacturer’s protocol. 1 μg of DNA construct in each case was used with 1.5 μL Lipofectamine 3000 Reagent. Media was changed within 24 hr post transfection and cells on the coverslips were collected 72 hr post transfection for indirect immunofluorescence staining.

For CasRx guide validation, HEK293T cells (ATCC, Manassas, VA) were seeded at 3x*10*^5^ cells per well in 6-well plates using Dulbecco Modified Eagle’s Medium (DMEM) containing L-glutamax (ThermoFisher Scientific, Rockford, IL) supplemented with 10% FBS (Gemini, West Sacramento, CA) 24 hr prior to transfection. Cells were transfected using Lipofectamine P3000 reagents (ThermoFisher Scientific, Rockford, IL) with 150 ng of each cDNA ORF plasmid containing a Myc-DDK tagged version of each target gene (Origene, Rockville, MD; Scn4b #MR226154) and 2350 ng of pAAV CasRx construct containing one guide targeting that gene. Media was removed and new media containing 4 μg/mL of puromycin was added for selection. 48 hr post puromycin selection, cells were collected for knockdown analysis via Western Blot and qPCR.

### Western blotting

HEK293T cells from the guide validation experiment were washed 3 times with PBS to remove excess media, resuspended in RIPA lysis buffer containing an EDTA-free protease inhibitor cocktail (Millipore Sigma, Billerica, MA), lysed by passing through a 25-gauge needle 10 times, and incubated on ice for 20 min. Lysates were centrifuged at 22,000 x *g* for 15 min at 4°C and the resulting supernatant was collected. Bicinchoninic acid assay (BCA; ThermoFisher Scientific, Rockford, IL) measurements were performed using the diluted protein lysates to normalize the amount of total protein loaded on gels across different samples. Samples containing 10 μg of protein mixed with 1 x LDS loading dye (ThermoFisher Scientific, Rockford, IL) were denatured at 70°C for 10 min and loaded onto 4-12% Bis-Tris gels (ThermoFisher Scientific, Rockford, IL). The gels were run using a MES running buffer (ThermoFisher Scientific, Rockford, IL) at 175V for 1 hr. The protein samples were then transferred onto PVDF membranes using an iBlot dry transfer apparatus (ThermoFisher Scientific, Rockford, IL). Membranes were washed 3 times with 1XPBS containing 0.05% Tween20 (ThermoFisher Scientific, Rockford, IL), blocked with 5% milk for 1 hr at room temperature, and incubated with anti-FLAG primary antibody (Cell Signaling Technology, Danvers, MA, #14793, 1:1000) at 4°C overnight with gentle orbital shaking. Membranes were later washed 3 times with 1XPBS-T and incubated with secondary HRP (ThermoFisher Scientific, Rockford, IL; 1:10,000) for 1 hr at room temperature with gentle orbital shaking. Lastly, membranes were washed again 3 times with 1XPBS-T and developed using the Pierce ECL Plus substrate (ThermoFisher Scientific, Rockford, IL) for chemiluminescent detection. ImageJ software was used to quantify the intensity of the bands to determine the overall knockdown levels.

### Quantitative real-time PCR (qRT-PCR) to assess *Scn4b* KD

RNA was isolated from HEK293T cells from the guide validation experiments using the RNeasy Mini Kit (Qiagen, Hilden, Germany) or from the heart of *Scn4b* KD mice using the AllPrep DNA/RNA/Protein Kit (Qiagen, Hilden, Germany). 1 μg of the resulting RNA was used for cDNA synthesis using QuantiTect Reverse Transcription Kit (Qiagen, Hilden Germany). For qRT-PCR, SYBR Green Master Mix (ThermoFisher Scientific, Rockford, IL) was used with the following primer pairs:

***Scn4b***: FWD - 5’tcctgaagaagacccgagag3’ REV - 5’ggcaacccgttctctgtg3’

***Actb***: validated mouse positive control primer sets (Active Motif, Carlsbad CA, #71015) PCR reactions in 96-well plates were run on a StepOnePlus system (ThermoFisher Scientific, Rockford, IL). The relative expression of the target genes was calculated based on the Ct values of *Scn4b* normalized to the Ct values of *Actb*.

### Sample harvest

All brain dissections were performed on ice after cooling the head in liquid nitrogen for 3 s. Dissected whole brain, hemi brains, and individual brain regions were subsequently flash frozen in liquid nitrogen and stored at −80 °C until further use. Striatal dissections were performed using a punch dissection method. In brief, mouse craniums were flash frozen in liquid nitrogen after decapitation. The brains were extracted, placed onto the pre-chilled Alto acrylic brain coronal mold (CellPoint Scientific, Gaithersburg, MD), and cut at positions #3 and #5 using razor blades. The resulting coronal slice was placed on a metal block on ice and the 2 mm biopsy punch (Miltex, York, PA) was used to collect tissues from the dorsal striatum. All brain perfusions for immunofluorescence and *in situ* experiments were performed using transcardial perfusion with 4% PFA in 1x PBS.

### Indirect immunofluorescence (IF) staining

For nuclear localization staining, cells on coverslips were washed 3 times with 1X PBS (ThermoFisher Scientific, Rockford, IL) to remove any remaining media. Cells were fixed in 4% paraformaldehyde (PFA; Electron Microscopy Solutions, Hatfield, PA) in 1X PBS solution for 1 hr at room temperature and washed 3 times with 1X PBS solution.

Next, cells were permeabilized with 1X PBS with 0.1% TritonX (PBS-T) for 15 min, washed 3 times with 1X PBS-T, and blocked with PBS-based blocking buffer containing 0.3M glycine (Sigma, St. Louis, MO) and 1% IgG-free BSA (Jackson Immunoresearch, West Grove, PA). Cells were then incubated with anti-HA primary antibody (Cell Signaling Technology, Danvers, MA; #3724, 1:100) at 4°C overnight with gentle orbital shaking, washed 3 times with PBS-T, and incubated in the AlexaFluor secondary antibody (ThermoFisher Scientific, Rockford, IL, 1:500) for 1 hr at room temperature. Lastly, cells were washed again 3 times with PBS-T, stained with DAPI, and mounted with ProLong Gold antifade mounting medium (ThermoFisher Scientific, Rockford, IL, 1:500). Cells were imaged with a Leica Stellaris Microscope to determine the nuclear localization pattern of HA-CasRx.

For mouse brain tissue staining, paraformaldehyde-perfused mouse brain sections stored at −80°C were washed with 1X tris-buffer saline solution (TBS; ThermoFisher Scientific, Rockford, IL), permeabilized with 0.2% Triton X-100 in 1X PBS for 15 min at RT and blocked with TBS-based blocking buffer containing 2% heat-inactivated donkey serum, 0.1% fish gelatin in TBS with 0.1% Triton X-100 for 1 hr at RT. Samples were then incubated with anti-HA (Cell Signaling Technology, Danvers, MA, #3724, 1:100), anti-NeuN (Synaptic Systems, Goettingen, Germany, #266004, 1:500), anti-IBA1 (FUJIFILM Wako Pure Chemical Corporation, #019-19741, 1:500) or anti-GFAP (Abcam, Waltham, MA, 1:2000) primary antibody at 4°C overnight with gentle orbital shaking. Following three 1XTBS washes, samples were incubated with AlexaFluor-conjugated secondary antibodies (ThermoFisher Scientific, Rockford, IL, 1:500) for 1 hr at RT with gentle orbital shaking, washed three times in 1XTBS, and counterstained with DAPI. Lastly, samples were mounted with ProLong Gold antifade mounting medium (ThermoFisher Scientific, Rockford, IL, 1:500). Whole sagittal sections were imaged on a Leica Stellaris Microscope and multiple striatal sections were analyzed for transduction efficiency and immune activation signal using CellProfiler.

### Combined IF and RNA fluorescence *in situ* hybridization (IF - RNA FISH)

Combined IF-RNA FISH was performed following the manufacturer’s technical note for Integrated Co-Detection Workflow (ICW) and the Multiplex Fluorescent Reagent Kit v2 (Advanced Cell Diagnostics, Newark, CA, #323100) using the manufacturer’s protocol for fresh-frozen tissues. Paraformaldehyde-perfused mouse brain sections stored at - 80°C were post-fixed in 4% paraformaldehyde (PFA; Electron Microscopy Solutions, Hatfield, PA, #15714) in 1X PBS for 1 hr at 4°C and dehydrated through an ethanol series of 50%, 70%, and 100%. A hydrophobic barrier was created around the sections using the ImmEdge hydrophobic barrier pen (Vector Laboratories, Newark, CA, #101098-065). Samples were then treated with hydrogen peroxide, permeabilized and blocked as described in **Indirect immunofluorescence (IF) staining** and then incubated with anti-HA (Cell Signaling Technology, Danvers MA, #3724, 1:100), anti-Bcl11b (Abcam, Waltham MA, #ab18465, 1:200), or anti-EM48 (Millipore Sigma, Billerica MA, #MAB5374, 1:200) primary antibodies in Co-Detection Anitibody Dilutent from the RNA-Protein Co-Dectction Ancillary kit (Advanced Cell Diagnostics, Newark, CA, #323180) overnight at 4°C with gentle orbital shaking. Samples were then washed 3 times in 1XTBS with 0.1% Triton X-100, fixed in 4% PFA for 30 min at RT and treated with Protease Plus for 30 min at 40°C. Next, samples were incubated with RNAScope probe targeting Ms-*SCN4B* (#484771; Advanced Cell Diagnostics, Newark, CA) for 2 hr at 40°C. Samples were then submerged in 5X saline-sodium citrate solution overnight at RT. Signals were amplified through 3 amplification steps, developed using the Opal dyes (Opal 570 #FP1488001KT; Akoya Biosciences, Marlborough, MA) at 1:3000 dilution. After the Opal dyes developing step, samples were then incubated in AlexaFluor-conjugated secondary antibodies, counterstained with DAPI, and mounted with ProLong Gold antifade mounting medium (ThermoFisher Scientific, Rockford, IL).

For human post-mortem caudate nucleus samples, a similar protocol was followed with additional pre-treatment steps. In short, following PFA fixation, samples were photobleached using a previously published protocol^40^ by which samples were exposed to a 300 W full-spectrum LED light source (PlatinumLED Therapy Lights, Carrollwood, FL, P300) while submerged in cold 1XPBS overnight to quench lipofuscin autofluorescence before the dehydration steps. Additionally, samples were treated with freshly diluted TrueBlack Lipofuscin Autofluorescence Quencher (Biotium, Fremont, CA, #23007, 1:20 in 70% ethanol) for 10 s before mounting.

### Adeno-associated virus (AAV) production and quantification

AAV PHP.eB virus was produced in-house following the protocol outlined in a published protocol.^41^ In brief, HEK293T/17 cells were cultured as described above and seeded at 5×10^6^ cells per 60-mm tissue culture dish (Corning, Glendale, AZ) in 5% FBS 24 hr prior to transfection. Cells were triple transfected with pAAV constructs, pHelper (Agilent), and pUCmini-iCAP-PHP, a gift from Viviana Gradinaru (Addgene plasmid #103005) using polyethyleneimine (PEI) at 90% confluency. At 24 hr post transfection, the media was discarded and replaced with fresh media. At 72 hr post transfection, the media containing viral particles was collected (and stored at 4°C) and replaced with fresh media. At 120 hr post transfection, both media containing viral particles and cells were harvested, combined with the media collected at 72 hr, and centrifuged at 2,000 x *g* for 15 min at RT. The supernatant was collected and mixed with 40% PEG (wt/vol; ThermoFisher Scientific) at 4°C overnight. The cell pellet was resuspended in salt-active nuclease (SAN) solution containing 100U/mL SAN (ArcticZymes, Tromsø, Norway) and incubated at 37°C for 1 hr before transferring to 4°C overnight. The PEG mixture was centrifuged at 4,000 x *g* for 30 min at 4°C, resuspended in SAN solution containing 100U/mL SAN, combined with the cell lysate fraction, and incubated at 37°C for 30 min. AAVs were purified by on equilibrium zonal centrifugation on iodixanol gradients (15%, 25%, 40%, and 60%), from which the purified virus was collected from the 40/60% interface and buffer-exchanged 3 times into 1XPBS using a 100 kDa Amicon filter device (Millipore Sigma, St. Louis, MO). Finally, a Costar Spin-X centrifuge tube filter (Corning, Glendale, AZ) was used to filter-sterilize the virus.

Viral titers were quantified using the AAVpro Titration Kit (for Real Time PCR, Ver. 2; Takara Bio USA, San Jose, CA) with AAVhSyn-GFP virus obtained from Virovek (Houston, TX) and SignaGen Laboratories (Frederick, MD) serving as internal controls. Purity assessment was done as previously described^42^ using silver protein stains (ThermoFisher Scientific, Rockford, IL) to detect the three viral protein bands of VP1, VP2, and VP3.

### Retro-orbital sinus intravenous AAV injection

Mice were anesthetized with 2% isoflurane in an induction chamber before being transferred to an anesthesia nose cone. One drop of 0.5% proparacaine (Bernell) was added to each eye before injection. 100uL of AAV virus containing either 1E12 vg or 5E12 vg was administered using a 27.5-gauge insulin needle (BD Biosciences, San Diego, CA) inserted at a 45-degree angle into the retro-orbital sinus. Optixcare Eye Lube (Med Vet International, Mettawa, IL) was then applied to both eyes. The mouse was then placed back into the homecage with the head resting on a nestlet and monitored until fully recovered.

### Behavioral testing

#### Open Field Test

Open field testing was performed as previously described.^43^ Mice were placed one at a time in the center of the sound-insulated open field box with 16 infrared beams spaced regularly along the x, y, and z axes (17” X 17” x 12”; Med Associates) and allowed to explore freely for the 30 min for CasRx-mediated Scn4b KD or 60 min for all other experiments. All experiments were performed during the light phase starting at 7AM and finishing before 1PM except for the zQ175 experiment mice, where the test was performed during dark phase starting at 7PM and finishing before 1AM. Mice from different experimental groups were tested simultaneously in a given run to ensure that all experimental groups were represented at the same phase of the testing time and to avoid time-of-day testing artifacts. Horizontal distance traveled, resting time, and ambulatory, vertical, and stereotypic activities (time and count) were recorded by the testing software.

#### Rotarod Test

Rotarod testing was performed as previously described^44^ using an accelerating rotarod apparatus (Med Associates, St. Albans, VT). Mice were trained on a rod that rotated at a constant speed of 20 rotations per min (RPM) for the duration of 5 min three times during the training phase on Day 1. They were repeatedly placed back on the rod once they fell and the total number of falls was recorded. After 2 hr of rest, during the testing phase, they were placed back on the rod that was now accelerating from 5 to 40 RPM over 5 min, three times. The duration that mice stayed on the rod until they fell or completed two or more passive rotations was recorded as latency to fall. The testing was repeated for 3 to 5 consecutive days without cage changing between sessions.

#### Rearing and Climbing Test

Rearing and climbing testing was performed following the published protocol.^45^ In brief, mice were placed one at a time underneath a wire mesh pencil cup (4.375” diameter x 5.5” height, Rolodex#82406) in front of a mirror for the duration of 5 min. Video recording was performed and used to count the number of rearing and climbing times for the duration of testing. Blinded scoring was performed using a randomized letter code assigned during testing.

#### Novel Object Recognition Test

The novel object recognition testing was performed following the published protocol.^46^ In brief, mice were placed one at a time in a sound-insulated open field box with 16 infrared beams spaced regularly along the x, y, and z axes (17” X 17” x 12”; #MED-OFAS-RSU, Med Associates, St. Albans, VT) and allowed to explore for the duration of 60 min similar to the open field test, but with the addition of two objects that were taped to the bottom of the box. These two objects were placed along the centerline of the box, each 1/6 of the box width from their corresponding walls. For the familiarization stage, two circular lids of a 15mL conical tube (ThermoFisher Scientific, Rockford, IL) were used as object one and object two. For the testing stage, one of the circular lids was replaced with a small binder clip (Uline, Pleasant Prairie, WI). Settings were used for the analysis such that the area of the box was subdivided into nine small squares and the ambulatory time for each given square was reported. The duration that mice explored the area containing the objects was used as the time exploring each object. The novel object discrimination index was calculated by dividing the difference in time exploring the new and the familiar object by the total time.

#### Burrowing Test

The burrowing assay was performed by placing one mouse per cage in a large cage containing a red rat tunnel (Bioserv, Prospect, CT) filled with 140g of regular mouse food pellets for a duration of 2 hr. The weight of the food remaining in the tube was measured at the end of the test and the percent of food pellet removed from the tube (burrowed) was calculated.

#### Marble Burying Test

The marble burying test was performed by placing one mouse per cage in a cage containing 5 cm in height of bedding material with 20 marbles equally spaced out in five rows and four columns for the duration of 30 min. The total number of marbles displaced from their original position was calculated and reported.

#### Gait Analysis

The gait analysis test was adapted from the published protocol.^47^ In brief, the four paws of each mouse were painted with different colors using non-toxic paint (Crayola, Easton, PA). The mice were then allowed to walk on the sheet of thick white Whatman filter paper (Cytiva, Marlborough, MA). The footprints were collected and analyzed for stride and stance lengths as well as fore/hind overlap. The data presented represents an average value from five consecutive steps for a given mouse.

#### Arousability Test

Arousability testing was conducted as previously described.^44^ The fraction of mice displaying responsive arousal to the cage lid opening was determined immediately after opening the lids of their home cages.

#### Righting Reflex

Mice were placed on their side and the duration that they required to right themselves and stand back in their normal position was recorded.

#### Hindlimb Clasping Test

The hindlimb clasping test was performed following the published protocol.^48^ In brief, mice were held by the base of their tail and lifted out of the cage for 10 sec. A video recording was performed and used for blinded scoring of the degree of dystonic clasping of the hind limbs.

#### Grip Strength

The grip strength assay was performed using a mouse grip strength meter, a T-shaped bar connected to a force-rate monitor (Ugo Basile, Varese, Italy). Specifically, mice were handled by the tail and gently placed onto the metal bar such that they held the bar with their forelimbs, and then they were gently pulled back until they released their grip. The test was repeated three times and the average value of the peak grip force in each trial was recorded.

#### Beam Walking Test

The beam walking test was performed using a specialized mouse beam walking apparatus (Maze Engineers, Skokie, IL). Specifically, mice were placed in a dark box located at the end of the beam along with their bedding material and food for 2 min during the familiarization stage. They were later placed on the opposite end of the beam and allowed to cross back into the dark box during the testing stage. The time required for them to successfully cross the beam was recorded. The test was repeated 3 times with 10 sec rest in between each run. A total of four different rod sizes were used with 10 min rest in between each rod.

#### Nest-building Test

The nest-building test was performed following the published protocol.^49^ In brief, mice were placed one at a time in a new cage containing bedding material, food pellets, water, and a pressed 5-cm cotton square nestlet (Ancare, Bellmore, NY) overnight starting from 7PM and until 9AM the next day. The weights of the nestlet before and after testing were measured and the percent nesting was calculated.

### Single Nuclear RNA Sequencing and Analysis^12^

#### Isolation of nuclei from fresh mouse brain tissue

Nuclei isolation was performed as reported in Lee et al. (Neuron 2020).^12^ All procedures were performed on ice. Briefly, each tissue was homogenized in 700 µL of homogenization buffer with a 2 mL tissue grinder using 10 strokes with a loose pestle, followed by 10 strokes with a tight pestle. Homogenized tissue was filtered through a 40 µm cell strainer and mixed with 450 µL of working solution. The mixture was then slowly pipetted onto the top of an iodixanol density gradient (300 µL 40% solution and 750 µL of 30% solution) inside a dolphin microcentrifuge tube. Nuclei were pelleted at the 30%-40% interface of the density gradient by centrifugation at 10,000 x *g* for 5 min at 4°C using a fixed angle rotor. The nuclear pellet was collected by aspirating ∼100µL from the interface and transferring to a 1.5 mL siliconized microcentrifuge tube. The pellet was washed with PBS with 2% BSA containing 0.12 U/µL SUPERase-In RNase Inhibitor (ThermoFisher Scientific, Rockford, IL). The nuclei were pelleted and washed by centrifugation at 300 x *g* for 3 min at 4°C using a swing-bucket rotor. The nuclei were again pelleted and washed twice more under the same conditions. The nuclear pellet was resuspended in 100 µL of PBS with 2% BSA.

#### Library preparation and sequencing

Droplet-based snRNA sequencing libraries were prepared using the Chromium Next GEM Single Cell 3′ reagent kit v3.1 and v4 (10x Genomics) according to the manufacturer’s protocol, targeting 10,000 nuclei per sample. Libraries were sequenced on a NovaSeq X at the Broad Institute Genomics Platform.

#### Sequencing data preprocessing

Cell Ranger v8.0 (10x Genomics) was used for genome alignment and feature-barcode matrix generation. Reads were aligned to the mouse reference genome mm10 allowing mapping to intronic regions.

#### Cell clustering and curation

We used the ACTIONet^50,51^ R package for data cleaning, batch correction, clustering, imputation, and cell type annotation as in Pineda et al. (Cell 2024)^37^ with some modifications. In brief, raw count matrices were prefiltered by removing barcodes containing less than 700 unique genes or greater than 20% mtRNA content. Counts were depth-normalized across samples and log-transformed. Further per-cluster filtering was performed by iteratively removing cells with abnormally low or high RNA content (relative to the distribution of the specific cluster), ambiguous overlapping profiles resembling dissimilar cell types (generally corresponding to doublet nuclei), and cells corresponding to graph nodes with a low *k*-core or low centrality in the cell-cell similarity network (generally corresponding to high ambient RNA content or doublet nuclei).

#### Cell type annotation

A curated set of known major cell type markers derived from Lee et al. (Neuron 2020)^12^ and Matsushima et al. (Nat Comm 2023)^28^ was used to coarsely annotate individual cells with their expected cell type and assign a confidence score to each annotation.

Large heterogeneous clusters were further partitioned based on visually distinct, well-defined gene expression domains reproducibly identified by Leiden^52^ and archetypal analysis^53^ clustering at multiple resolutions.

#### Differential gene expression analysis

We employed a pseudo-bulk approach considering each mouse as a single biological replicate for each genotype-condition pair per cell type. In some cases, small subclusters that lacked sufficient representation to be analyzed independently were merged at a higher annotation level, provided they all belonged to the same major cell type. We required that a cluster be represented across at least 3 mice per test group, separately for each experiment, and with at least 30 cells per replicate. If this criterion could not be met, these clusters were excluded from analysis. For each differential expression cluster, only genes present in at least 10% of cells were retained for analysis. Counts were depth normalized, scaled to 10,000 (target UMI count), and log_2_-transformed per cell. Pseudo-bulk expression profiles were computed by averaging normalized log-counts within each cluster for each unique sample. We used surrogate variable analysis^54^ (implemented in the SVA R package) to identify and remove sources of unknown variance, which were mostly negligible. Pseudo-bulk differential gene expression analysis was performed using the limma R package^55^ using the inverse of the per-sample single cell-level variance as weights. Genes were considered differentially expressed (DEGs) if they had a log_2_-fold change > or = to 1 standard deviation [fold change abs(*z*) >= 1] and with an FDR-corrected *p* adjusted value < 0.05, similar to previously reported.^37^

#### Pathway enrichment analysis

Gene Ontology Biological Process (GO:BP) pathway enrichment analysis was performed using the enrichR package. For each cell type and contrast, significant differentially expressed genes (DEGs) were defined using an adjusted p-value threshold of < 0.05. Enrichment analysis was conducted against the *GO_Biological_Process_2025* database. For visualization, pathways were ranked by adjusted p-value and the top pathways (Top 10) were selected per cell type and direction, and statistical significance was displayed as −log₁₀(adjusted p-value).

### Electrophysiological recordings

#### Slice preparation

Mice aged 15-18 months from the *Scn4b* OX study were used for the electrophysiological recording. After animals were anesthetized with isoflurane, brain tissue was quickly removed following decapitation in ice-cold solution containing (in mM) 90 sucrose, 80 NaCl, 3.5 KCl, 0.5 CaCl_2_, 4.5 MgCl_2_, 24 NaHCO_3_, 1.25 NaH_2_PO_4_, and 10 glucose saturated with 95% O_2_ and 5% CO_2_. Coronal slices (240 µm) were prepared using a vibratome (Leica) and kept in warm (∼32℃) artificial cerebrospinal fluid (ACSF) containing (in mM) 124 NaCl, 2.5 KCl, 2.5 CaCl_2_, 1.3 MgCl_2_, 26.2 NaHCO_3_, 1 NaH_2_PO_4_, 20 glucose, and 0.4 ascorbic acid saturated with 95% O_2_ and 5% CO_2_ for 20 min and then returned to room temperature until recording. For recording, slices were transferred to a submerged recording chamber and perfused at 2 ml/min with ACSF saturated with 95% O_2_ and 5% CO_2_, which was heated at 32℃ using an inline heater (Harvard Apparatus, Holliston, MA).

#### Slice electrophysiology

Whole-cell current clamp recordings were conducted in the dorsal striatum. The recordings were made from GFP-positive cells in wild-type and zQ175 mice, or from unlabeled cells with a soma diameter of ∼15 µm in the animals injected with the *Scn4b* OX viral vector at 15-18 months of age (12-15 months after injection). Recordings were made using glass electrodes (2.5–3.5 MΩ) filled with an internal solution containing (in mM): 120 K-methanesulfate, 20 KCl, 2 MgCl_2_, 10 HEPES, 0.2 K-EGTA, 4 Na-ATP, and 0.4 Na-GTP (pH ∼ 7.3, ∼290 mOsm). After achieving whole-cell configuration, the resting membrane potential was measured in current-clamped mode without current injection (with I=0). Typical SPNs showed the resting potentials around −90 mV. Then, cells were held around −80 mV with bias current and current steps (duration = 1 sec; amplitude = −50pA to 500 pA, in 50 pA increments) were delivered to elicit action potentials. Voltage signals from Multiclamp 700B (Molecular Devices) were filtered at 1 kHz and acquired at 5 kHz using Digidata 1550B and pClamp software (Molecular Devices, San Jose, CA). The AP thresholds were determined as the voltage at which the slope of the action potential reached 20 mV/ms.^56^ All data were analyzed using Igor Pro 7 (Sutter Instruments). Statistical analysis was carried out in Prism10.1 (GraphPad, San Diego, CA).

## QUANTIFICATION AND STATISTICAL ANALYSIS

### Cell Profiler Method for Transduction Efficiencies Analysis

To determine the viral transduction efficiency, we determined the number of neurons that were positive for HA tag signal. Whole striatum of mouse brain sagittal sections stained for HA, NeuN, and DAPI via indirect immunofluorescence staining were imaged at 40x magnification, each recorded with a section of 2048×2048 pixels. Within the CellProfiler framework, the images were loaded as grayscale. The identification of neurons was performed by overlapping the NeuN and DAPI signals as described above. After identifying neurons, we used the “MeasureObjectIntensity” module to measure the intensity of HA-stained images, loaded without any correction, within the area defined as neurons above. The module records the HA signal’s integrated intensity within a given object. In the post-processing step, we calculated the median HA-integrated intensity within each image and classified neurons with HA-integrated intensity greater than the median as HA-positive neurons. The transduction efficiency of each image was then calculated as HA+ divided by the total number of neurons within a given image.

### Cell Profiler Method for Immune Activation of IBA1 and GFAP Quantification

The IBA1 and GFAP intensity analyses were done on a per-image basis. In short, we used the “MeasureImageIntensity” module to measure the “TotalIntensity” of IBA1 or GFAP and NeuN signals within a given image. Dividing this total sum intensity of IBA1 or GFAP by that of NeuN gave the normalized sum intensity. In the post-process analysis, the intensities were normalized again, setting the mean of the IBA1 or GFAP Normalized Sum Intensity of the PBS sample to 1.

To count IBA cells, we enhanced the detection of the IBA signal using the “EnhanceOrSuppressFeatures” module by enhancing the “Texure” feature with a Smoothing scale of 5 after correcting the IBA1 images with “CorrectIllumination Calculate” and “CorrectIlluminationApply”. This procedure concentrated the IBA1 signal, making it easier to identify. We then used the “IndentifyPrimaryObjects” to identify the IBA1 puncta on a per-image basis. The IBA1 count within a given image was recorded.

Given the complexity of the GFAP images, we performed manual counting for the GFAP count based on the cell morphology. Cells were considered GFAP positive if the DAPI signal was localized to the center of the GFAP signal with the extended processes. The total number of GFAP counts within a given image was recorded.

### Cell Profiler Method for Scn4b Puncta and Intensity Quantification

A CellProfiler pipeline was used to quantify Scn4b puncta number from 63x magnification images, each with a section of 2048×2048 pixels. The DAPI objects, representing cell nuclei, were identified using the “IdentifyPrimaryObjects” module, followed by the identification of neurons (with NeuN or Bcl11b) using the “IdentifyPrimaryObjects” module. Then, the “RelateObjects” module was used to filter out the spurious detection of NeuN or Bcl11b signal by matching DAPI as parent objects with neuron-identifying objects. Since DAPI and the NeuN stains may not fully cover the entire neuron, making downstream quantification such as intensity measurement and puncta count uncertain, the size of the neuron objects needs to be enlarged, with different degrees depending on the experiment. This was done by applying the “ExpandOrShrinkObjects” module to the DAPI objects. The puncta were identified using the “IdentifyPrimaryObjects” module with adaptive Otsu method, 3px smoothing filter, and 5-40px size filter. The total number of puncta per neuron was reported. For the intensity quantification, “MeasureObjectIntensity” module, recording the “IntegratedSumIntensity” was used on the uncorrected images. The normalized SumIntensity of *Scn4b* and Bcl11b or NeuN was reported.

### Cell Profiler Method for EM48 and P90 Quantification

The analysis of the mHTT puncta was performed using CellProfiler pipeline on 63X magnification images. The channels stained for DAPI, NeuN, and EM48 or P90 were corrected using the “CorrectIllumination Calculate” and “CorrectIlluminationApply” modules in the respective order. For a given image set, cells were identified by first applying the “IdentifyPrimaryObjects” module to the DAPI channel using the adaptive Otsu method, 5px smoothing filter, and 15-90px size cutoff. Then, NeuN objects were identified using the “IdentifyPrimaryObjects” module with the global Otsu method, automatic smoothing filter, and 30-120px size cutoff. Neurons were identified by overlapping the DAPI objects with NeuN objects using the “RelateObjects” module, with DAPI as parent objects. EM48 and P90 objects were identified using the “IdentifyPrimaryObjects” module with adaptive Otsu method, 3px smoothing filter, and 5-40px size cutoff. The normalized EM48 and P90 count was calculated on a per-image basis by dividing the total count by the number of neurons.

## ACKNOWLEDGEMENTS

This research was supported by NIH/NINDS award R35 NS127327 (M.H.), intramural NIMH funding award ZIA MH002987 (V.A.A.), a fellowship from the Royal Thai Government (S.S.), and postdoctoral fellowship F32NS128067 from NIH/NINDS (R.L.). We thank tissue donors and their families, the NIH NeuroBioBank, the Netherlands Brain Bank, the University of Alabama at Birmingham, and the Harvard Brain Tissue Resource Center.

## AUTHOR CONTRIBUTIONS

S.S. – Conceptualization, Investigation, Formal Analysis, Writing – original draft

V.F. – Investigation, Writing – review & editing

S.P. – Investigation, Formal Analysis, Writing – original draft

H.L. - Investigation, Formal Analysis, Writing – review & editing

J.H.S. – Investigation, Formal Analysis, Writing – original draft

F.G. – Investigation, Writing – review & editing

R.M.L. – Investigation, Writing – review & editing

M.K. – Writing – review & editing, Resources, Supervision

V.A.A – Conceptualization, Formal Analysis, Writing – review & editing, Resources, Supervision

M.H. – Conceptualization, Formal Analysis, Writing – original draft, review & editing, Resources, Supervision

## DECLARATION OF INTERESTS

The authors declare no competing interests.

## Notes

### Competing Interest Statement

The authors have declared no competing interest.

## REFERENCES

1. Reiner, A., Albin, R.L., Anderson, K.D., D’Amato, C.J., Penney, J.B., and Young, A.B. (1988). Differential loss of striatal projection neurons in Huntington disease. Proc. Natl. Acad. Sci. U. S. A. 85, 5733–5737. 10.1073/pnas.85.15.5733.

2. Ross, C.A., Aylward, E.H., Wild, E.J., Langbehn, D.R., Long, J.D., Warner, J.H., Scahill, R.I., Leavitt, B.R., Stout, J.C., Paulsen, J.S., et al. (2014). Huntington disease: natural history, biomarkers and prospects for therapeutics. Nat. Rev. Neurol. 10, 204–216. 10.1038/nrneurol.2014.24.

3. A novel gene containing a trinucleotide repeat that is expanded and unstable on Huntington’s disease chromosomes. The Huntington’s Disease Collaborative Research Group (1993). Cell 72, 971–983. 10.1016/0092-8674(93)90585-e.

4. Hong, E.P., MacDonald, M.E., Wheeler, V.C., Jones, L., Holmans, P., Orth, M., Monckton, D.G., Long, J.D., Kwak, S., Gusella, J.F., et al. (2021). Huntington’s Disease Pathogenesis: Two Sequential Components. J. Huntingt. Dis. 10, 35–51. 10.3233/JHD-200427.

5. Wang, N., Zhang, S., Langfelder, P., Ramanathan, L., Gao, F., Plascencia, M., Vaca, R., Gu, X., Deng, L., Dionisio, L.E., et al. (2025). Distinct mismatch-repair complex genes set neuronal CAG-repeat expansion rate to drive selective pathogenesis in HD mice. Cell 188, 1524–1544.e22. 10.1016/j.cell.2025.01.031.

6. Mätlik, K., Baffuto, M., Kus, L., Deshmukh, A.L., Davis, D.A., Paul, M.R., Carroll, T.S., Caron, M.-C., Masson, J.-Y., Pearson, C.E., et al. (2024). Cell-type-specific CAG repeat expansions and toxicity of mutant Huntingtin in human striatum and cerebellum. Nat. Genet. 56, 383–394. 10.1038/s41588-024-01653-6.

7. Lee, J.-M., Huang, Y., Orth, M., Gillis, T., Siciliano, J., Hong, E., Mysore, J.S., Lucente, D., Wheeler, V.C., Seong, I.S., et al. (2022). Genetic modifiers of Huntington disease differentially influence motor and cognitive domains. Am. J. Hum. Genet. 109, 885–899. 10.1016/j.ajhg.2022.03.004.

8. Jimenez-Sanchez, M., Licitra, F., Underwood, B.R., and Rubinsztein, D.C. (2017). Huntington’s Disease: Mechanisms of Pathogenesis and Therapeutic Strategies. Cold Spring Harb. Perspect. Med. 7, a024240. 10.1101/cshperspect.a024240.

9. Genetic Modifiers of Huntington’s Disease (GeM-HD) Consortium (2025). Genetic modifiers of somatic expansion and clinical phenotypes in Huntington’s disease highlight shared and tissue-specific effects. Nat. Genet. 57, 1426–1436. 10.1038/s41588-025-02191-5.

10. Genetic Modifiers of Huntington’s Disease (GeM-HD) Consortium. Electronic address: and Genetic Modifiers of Huntington’s Disease (GeM-HD) Consortium (2019). CAG Repeat Not Polyglutamine Length Determines Timing of Huntington’s Disease Onset. Cell 178, 887–900.e14. 10.1016/j.cell.2019.06.036.

11. Baide-Mairena, H., Marti-Sánchez, L., Marcé-Grau, A., Cazurro-Gutiérrez, A., Sanchez-Montanez, A., Delgado, I., Moreno-Galdó, A., Macaya-Ruiz, A., García-Arumí, E., Pérez-Dueñas, B., et al. (2022). Genetic diagnosis of basal ganglia disease in childhood. Dev. Med. Child Neurol. 64, 743–752. 10.1111/dmcn.15125.

12. Lee, H., Fenster, R.J., Pineda, S.S., Gibbs, W.S., Mohammadi, S., Davila-Velderrain, J., Garcia, F.J., Therrien, M., Novis, H.S., Gao, F., et al. (2020). Cell Type-Specific Transcriptomics Reveals that Mutant Huntingtin Leads to Mitochondrial RNA Release and Neuronal Innate Immune Activation. Neuron 107, 891–908.e8. 10.1016/j.neuron.2020.06.021.

13. Langfelder, P., Cantle, J.P., Chatzopoulou, D., Wang, N., Gao, F., Al-Ramahi, I., Lu, X.-H., Ramos, E.M., El-Zein, K., Zhao, Y., et al. (2016). Integrated genomics and proteomics define huntingtin CAG length-dependent networks in mice. Nat. Neurosci. 19, 623–633. 10.1038/nn.4256.

14. Yu, F.H., Westenbroek, R.E., Silos-Santiago, I., McCormick, K.A., Lawson, D., Ge, P., Ferriera, H., Lilly, J., DiStefano, P.S., Catterall, W.A., et al. (2003). Sodium channel beta4, a new disulfide-linked auxiliary subunit with similarity to beta2. J. Neurosci. Off. J. Soc. Neurosci. 23, 7577–7585. 10.1523/JNEUROSCI.23-20-07577.2003.

15. Brackenbury, W.J., and Isom, L.L. (2011). Na Channel β Subunits: Overachievers of the Ion Channel Family. Front. Pharmacol. 2, 53. 10.3389/fphar.2011.00053.

16. Ji, X., Saha, S., Gao, G., Lasek, A.W., Homanics, G.E., Guildford, M., Tapper, A.R., and Martin, G.E. (2017). The Sodium Channel β4 Auxiliary Subunit Selectively Controls Long-Term Depression in Core Nucleus Accumbens Medium Spiny Neurons. Front. Cell. Neurosci. 11, 17. 10.3389/fncel.2017.00017.

17. Ransdell, J.L., Dranoff, E., Lau, B., Lo, W.-L., Donermeyer, D.L., Allen, P.M., and Nerbonne, J.M. (2017). Loss of Navβ4-Mediated Regulation of Sodium Currents in Adult Purkinje Neurons Disrupts Firing and Impairs Motor Coordination and Balance. Cell Rep. 19, 532–544. 10.1016/j.celrep.2017.03.068.

18. Oyama, F., Miyazaki, H., Sakamoto, N., Becquet, C., Machida, Y., Kaneko, K., Uchikawa, C., Suzuki, T., Kurosawa, M., Ikeda, T., et al. (2006). Sodium channel beta4 subunit: down-regulation and possible involvement in neuritic degeneration in Huntington’s disease transgenic mice. J. Neurochem. 98, 518–529. 10.1111/j.1471-4159.2006.03893.x.

19. Miyazaki, H., Oyama, F., Inoue, R., Aosaki, T., Abe, T., Kiyonari, H., Kino, Y., Kurosawa, M., Shimizu, J., Ogiwara, I., et al. (2014). Singular localization of sodium channel β4 subunit in unmyelinated fibres and its role in the striatum. Nat. Commun. 5, 5525. 10.1038/ncomms6525.

20. Blednov, Y.A., Bajo, M., Roberts, A.J., Da Costa, A.J., Black, M., Edmunds, S., Mayfield, J., Roberto, M., Homanics, G.E., Lasek, A.W., et al. (2019). Scn4b regulates the hypnotic effects of ethanol and other sedative drugs. Genes Brain Behav. 18, e12562. 10.1111/gbb.12562.

21. Konermann, S., Lotfy, P., Brideau, N.J., Oki, J., Shokhirev, M.N., and Hsu, P.D. (2018). Transcriptome Engineering with RNA-Targeting Type VI-D CRISPR Effectors. Cell 173, 665–676.e14. 10.1016/j.cell.2018.02.033.

22. Chan, K.Y., Jang, M.J., Yoo, B.B., Greenbaum, A., Ravi, N., Wu, W.-L., Sánchez-Guardado, L., Lois, C., Mazmanian, S.K., Deverman, B.E., et al. (2017). Engineered AAVs for efficient noninvasive gene delivery to the central and peripheral nervous systems. Nat. Neurosci. 20, 1172–1179. 10.1038/nn.4593.

23. Choi, J.-H., Yu, N.-K., Baek, G.-C., Bakes, J., Seo, D., Nam, H.J., Baek, S.H., Lim, C.-S., Lee, Y.-S., and Kaang, B.-K. (2014). Optimization of AAV expression cassettes to improve packaging capacity and transgene expression in neurons. Mol. Brain 7, 17. 10.1186/1756-6606-7-17.

24. Carter, R.J., Lione, L.A., Humby, T., Mangiarini, L., Mahal, A., Bates, G.P., Dunnett, S.B., and Morton, A.J. (1999). Characterization of progressive motor deficits in mice transgenic for the human Huntington’s disease mutation. J. Neurosci. Off. J. Soc. Neurosci. 19, 3248–3257. 10.1523/JNEUROSCI.19-08-03248.1999.

25. Medeiros-Domingo, A., Kaku, T., Tester, D.J., Iturralde-Torres, P., Itty, A., Ye, B., Valdivia, C., Ueda, K., Canizales-Quinteros, S., Tusié-Luna, M.T., et al. (2007). SCN4B-encoded sodium channel beta4 subunit in congenital long-QT syndrome. Circulation 116, 134–142. 10.1161/CIRCULATIONAHA.106.659086.

26. Jackson, K.L., Dayton, R.D., Deverman, B.E., and Klein, R.L. (2016). Better Targeting, Better Efficiency for Wide-Scale Neuronal Transduction with the Synapsin Promoter and AAV-PHP.B. Front. Mol. Neurosci. 9, 116. 10.3389/fnmol.2016.00116.

27. Ji, X., Saha, S., Gao, G., Lasek, A.W., Homanics, G.E., Guildford, M., Tapper, A.R., and Martin, G.E. (2017). The Sodium Channel β4 Auxiliary Subunit Selectively Controls Long-Term Depression in Core Nucleus Accumbens Medium Spiny Neurons. Front. Cell. Neurosci. 11, 17. 10.3389/fncel.2017.00017.

28. Matsushima, A., Pineda, S.S., Crittenden, J.R., Lee, H., Galani, K., Mantero, J., Tombaugh, G., Kellis, M., Heiman, M., and Graybiel, A.M. (2023). Transcriptional vulnerabilities of striatal neurons in human and rodent models of Huntington’s disease. Nat. Commun. 14, 282. 10.1038/s41467-022-35752-x.

29. Obenauer, J.C., Chen, J., Andreeva, V., Aaronson, J.S., Lee, R., Caricasole, A., and Rosinski, J. (2022). Expression analysis of Huntington disease mouse models reveals robust striatum disease signatures. Preprint at Neuroscience, 10.1101/2022.02.04.479180.

30. de Leeuw, C.N., Korecki, A.J., Berry, G.E., Hickmott, J.W., Lam, S.L., Lengyell, T.C., Bonaguro, R.J., Borretta, L.J., Chopra, V., Chou, A.Y., et al. (2016). rAAV-compatible MiniPromoters for restricted expression in the brain and eye. Mol. Brain 9, 52. 10.1186/s13041-016-0232-4.

31. Deng, Y., Joni, M., Wang, H., Cox, R., Sapp, E., DiFiglia, M., and Reiner, A. (2026). Localization of mutant huntingtin with HTT Exon1 P90 C-terminal neoepitope antibodies in relation to regional and neuronal vulnerability in forebrain in Q175 mice and human huntington’s disease. J. Huntingt. Dis. 15, 55–94. 10.1177/18796397251404999.

32. Cummings, D.M., Cepeda, C., and Levine, M.S. (2010). Alterations in striatal synaptic transmission are consistent across genetic mouse models of Huntington’s disease. ASN Neuro 2, e00036. 10.1042/AN20100007.

33. Heikkinen, T., Lehtimäki, K., Vartiainen, N., Puoliväli, J., Hendricks, S.J., Glaser, J.R., Bradaia, A., Wadel, K., Touller, C., Kontkanen, O., et al. (2012). Characterization of neurophysiological and behavioral changes, MRI brain volumetry and 1H MRS in zQ175 knock-in mouse model of Huntington’s disease. PloS One 7, e50717. 10.1371/journal.pone.0050717.

34. Brulé, B., Alcalá-Vida, R., Penaud, N., Scuto, J., Mounier, C., Seguin, J., Khodaverdian, S.V., Cosquer, B., Birmelé, E., Le Gras, S., et al. (2025). Accelerated epigenetic aging in Huntington’s disease involves polycomb repressive complex 1. Nat. Commun. 16, 1550. 10.1038/s41467-025-56722-z.

35. Seong, I.S., Woda, J.M., Song, J.-J., Lloret, A., Abeyrathne, P.D., Woo, C.J., Gregory, G., Lee, J.-M., Wheeler, V.C., Walz, T., et al. (2010). Huntingtin facilitates polycomb repressive complex 2. Hum. Mol. Genet. 19, 573–583. 10.1093/hmg/ddp524.

36. von Schimmelmann, M., Feinberg, P.A., Sullivan, J.M., Ku, S.M., Badimon, A., Duff, M.K., Wang, Z., Lachmann, A., Dewell, S., Ma’ayan, A., et al. (2016). Polycomb repressive complex 2 (PRC2) silences genes responsible for neurodegeneration. Nat. Neurosci. 19, 1321–1330. 10.1038/nn.4360.

37. Pineda, S.S., Lee, H., Ulloa-Navas, M.J., Linville, R.M., Garcia, F.J., Galani, K., Engelberg-Cook, E., Castanedes, M.C., Fitzwalter, B.E., Pregent, L.J., et al. (2024). Single-cell dissection of the human motor and prefrontal cortices in ALS and FTLD. Cell 187, 1971–1989.e16. 10.1016/j.cell.2024.02.031.

38. Spratt, P.W.E., Ben-Shalom, R., Keeshen, C.M., Burke, K.J., Clarkson, R.L., Sanders, S.J., and Bender, K.J. (2019). The Autism-Associated Gene Scn2a Contributes to Dendritic Excitability and Synaptic Function in the Prefrontal Cortex. Neuron 103, 673–685.e5. 10.1016/j.neuron.2019.05.037.

39. Labun, K., Montague, T.G., Krause, M., Torres Cleuren, Y.N., Tjeldnes, H., and Valen, E. (2019). CHOPCHOP v3: expanding the CRISPR web toolbox beyond genome editing. Nucleic Acids Res. 47, W171–W174. 10.1093/nar/gkz365.

40. Sun, Y., Ip, P., and Chakrabartty, A. (2017). Simple Elimination of Background Fluorescence in Formalin-Fixed Human Brain Tissue for Immunofluorescence Microscopy. J. Vis. Exp. JoVE, 56188. 10.3791/56188.

41. Challis, R.C., Ravindra Kumar, S., Chan, K.Y., Challis, C., Beadle, K., Jang, M.J., Kim, H.M., Rajendran, P.S., Tompkins, J.D., Shivkumar, K., et al. (2019). Systemic AAV vectors for widespread and targeted gene delivery in rodents. Nat. Protoc. 14, 379–414. 10.1038/s41596-018-0097-3.

42. Wörner, T.P., Bennett, A., Habka, S., Snijder, J., Friese, O., Powers, T., Agbandje-McKenna, M., and Heck, A.J.R. (2021). Adeno-associated virus capsid assembly is divergent and stochastic. Nat. Commun. 12, 1642. 10.1038/s41467-021-21935-5.

43. Hachigian, L.J., Carmona, V., Fenster, R.J., Kulicke, R., Heilbut, A., Sittler, A., Pereira de Almeida, L., Mesirov, J.P., Gao, F., Kolaczyk, E.D., et al. (2017). Control of Huntington’s Disease-Associated Phenotypes by the Striatum-Enriched Transcription Factor Foxp2. Cell Rep. 21, 2688–2695. 10.1016/j.celrep.2017.11.018.

44. Wertz, M.H., Mitchem, M.R., Pineda, S.S., Hachigian, L.J., Lee, H., Lau, V., Powers, A., Kulicke, R., Madan, G.K., Colic, M., et al. (2020). Genome-wide In Vivo CNS Screening Identifies Genes that Modify CNS Neuronal Survival and mHTT Toxicity. Neuron 106, 76–89.e8. 10.1016/j.neuron.2020.01.004.

45. Menalled, L.B., Kudwa, A.E., Miller, S., Fitzpatrick, J., Watson-Johnson, J., Keating, N., Ruiz, M., Mushlin, R., Alosio, W., McConnell, K., et al. (2012). Comprehensive behavioral and molecular characterization of a new knock-in mouse model of Huntington’s disease: zQ175. PloS One 7, e49838. 10.1371/journal.pone.0049838.

46. Leger, M., Quiedeville, A., Bouet, V., Haelewyn, B., Boulouard, M., Schumann-Bard, P., and Freret, T. (2013). Object recognition test in mice. Nat. Protoc. 8, 2531–2537. 10.1038/nprot.2013.155.

47. Menalled, L.B.; E.-K., Bassem F.;. Patry, Monica; Suárez-Fariñas, Mayte; Orenstein, Samantha J.;. Zahasky, Benjamin; Leahy, Christina; Wheeler, Vanessa C.;. Yang, X. William; MacDonald, Marcy E.;. Morton, A. Jennifer; Bates, GP; Leeds, Janet M.;. Park, Larry; Howland, David; Signer, Ethan; Tobin, Allan J.;. Brunner, Daniela (2009). Systematic behavioral evaluation of Huntington’s disease transgenic and knock-in mouse models. Neurobiol. Dis. 35, 319–336. 10.1016/j.nbd.2009.05.007.

48. Guyenet, S.J., Furrer, S.A., Damian, V.M., Baughan, T.D., La Spada, A.R., and Garden, G.A. (2010). A simple composite phenotype scoring system for evaluating mouse models of cerebellar ataxia. J. Vis. Exp. JoVE, 1787. 10.3791/1787.

49. Deacon, R.M. (2006). Assessing nest building in mice. Nat. Protoc. 1, 1117–1119. 10.1038/nprot.2006.170.

50. Mohammadi, S., Ravindra, V., Gleich, D.F., and Grama, A. (2018). A geometric approach to characterize the functional identity of single cells. Nat. Commun. 9, 1516. 10.1038/s41467-018-03933-2.

51. Mohammadi, S., Davila-Velderrain, J., and Kellis, M. (2020). A multiresolution framework to characterize single-cell state landscapes. Nat. Commun. 11, 5399. 10.1038/s41467-020-18416-6.

52. Traag, V.A., Waltman, L., and van Eck, N.J. (2019). From Louvain to Leiden: guaranteeing well-connected communities. Sci. Rep. 9, 5233. 10.1038/s41598-019-41695-z.

53. Cutler, A., and Breiman, L. (1994). Archetypal Analysis. Technometrics 36, 338–347. 10.1080/00401706.1994.10485840.

54. Leek, J.T., Johnson, W.E., Parker, H.S., Jaffe, A.E., and Storey, J.D. (2012). The sva package for removing batch effects and other unwanted variation in high-throughput experiments. Bioinformatics 28, 882–883. 10.1093/bioinformatics/bts034.

55. Ritchie, M.E., Phipson, B., Wu, D., Hu, Y., Law, C.W., Shi, W., and Smyth, G.K. (2015). limma powers differential expression analyses for RNA-sequencing and microarray studies. Nucleic Acids Res. 43, e47. 10.1093/nar/gkv007.

56. Bekkers, J.M., and Delaney, A.J. (2001). Modulation of excitability by alpha-dendrotoxin-sensitive potassium channels in neocortical pyramidal neurons. J. Neurosci. Off. J. Soc. Neurosci. 21, 6553–6560. 10.1523/JNEUROSCI.21-17-06553.2001.

