## Supplemental Figures for "Scn4b Modulates Huntington’s Disease Phenotype Severity in vivo"

### SUPPLEMENTAL MATERIAL

Supplemental Figures S1-9

Supplemental References

### SUPPLEMENTAL FIGURES

### Supplemental Figure 1

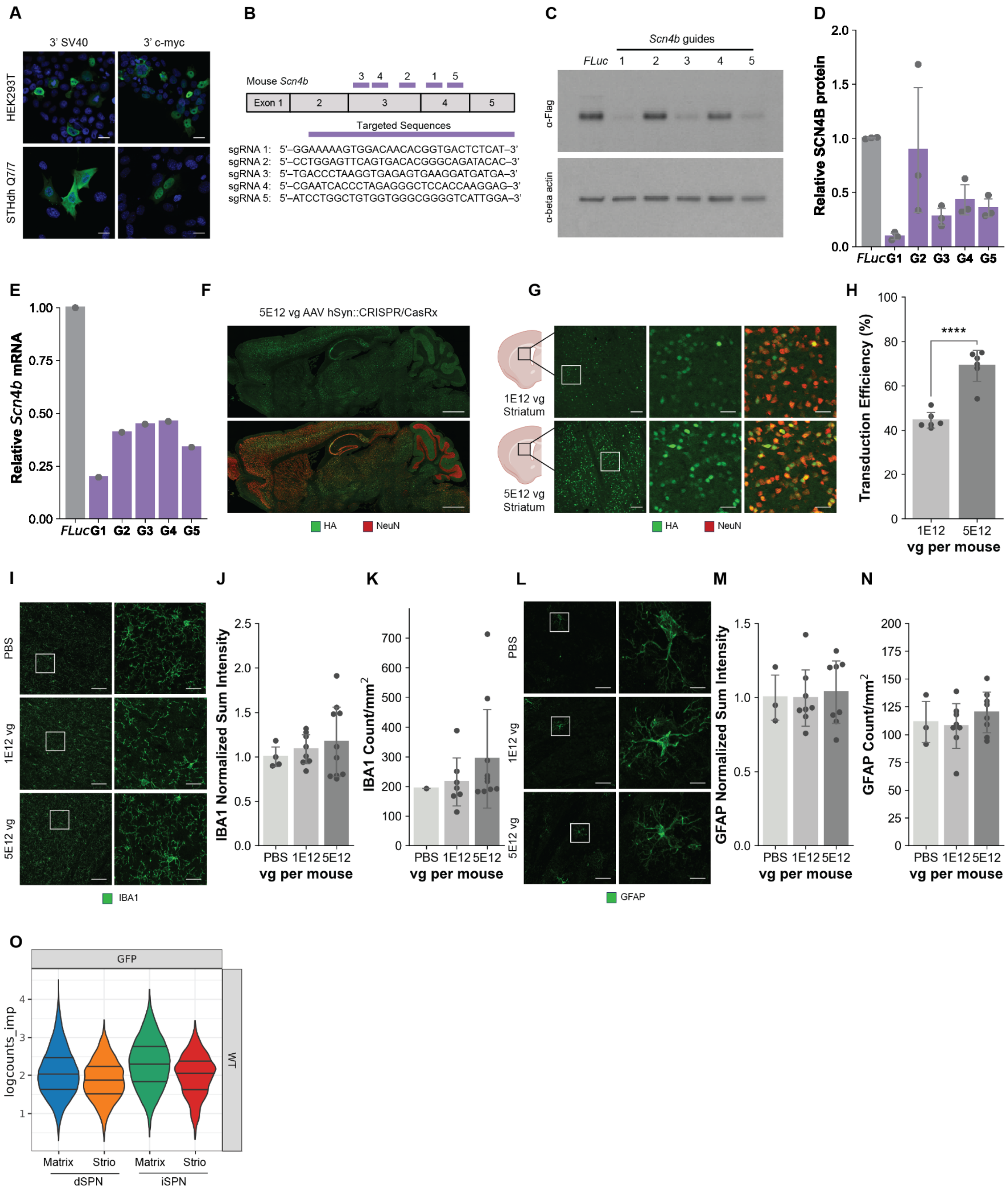

**Supplementary Figure 1. *In vitro* and *in vivo* optimization of AAV PHP.eB hSyn CRISPR/CasRx knockdown of *Scn4b*. (A)** Representative IF images demonstrating the CasRx localization pattern in HEK293T and STHdh Q7/7 cells transfected with an AAV construct containing CasRx flanked with either 5'SV40 NLS-CasRx-SV40 NLS 3' or 5'SV40 NLS-CasRx-bipartite cMyc NLS 3'. Green signal is the anti-HA stain for CasRx; blue signal is DAPI. Scale bar, 20 µm. **(B)** Guide design and schematic of guide binding along the exons of the *Scn4b* transcript. **(C)** Example Western blot analysis of guide knockdown (in these cell culture experiments, Scn4b is tagged with the FLAG epitope). **(D)** Quantification of *Scn4b* protein and **(E)** RNA knockdown level of each guide as determined by Western blot and qPCR, respectively. *FLuc* denotes the guides targeting Firefly Luciferase, which is not present in the mouse genome. Data are shown as mean ± SD. **(F)** Representative IF images of a sagittal brain section from a mouse injected with 5E12 vg of PHP.eB-hSyn::CRISPR/CasRx virus showing CasRx-3xHA (green) and NeuN (red). Scale bar, 1 mm. **(G)** Representative IF of striatal sections of mice injected with 5E12 vg and 1E12 vg of PHP.eB-hSyn::CRISPR/CasRx virus. Scale bar, 100 µm (left) and 25 µm (right). **(H)** Quantification of neuronal transduction efficiencies in the striatum. Data are shown as mean ± SD ($n$=6 mice per condition; 4 images (20x) per mouse). One-sided t-test: ****$p < 0.0001$. Representative IF staining for **(I)** IBA1 and **(L)** GFAP in mouse striatum injected with PBS, 1E12 vg, or 5E12 vg of AAV PHP.eB hSyn::CasRx-HA. Scale bar, 50 µm (left) and 10 µm (right). Quantification of fluorescence intensity and positive cell counts for IBA1**(J, K)** and GFAP **(M, N)**. Data are shown as mean ± SD. **(O)** Violin plot showing GFP expression (as reported by snRNA-seq data) driven by AAV PHP.eB under the hSyn promoter across striatal SPN subtypes showing higher expression in matrix than striosome SPNs.

### Supplemental Figure 2

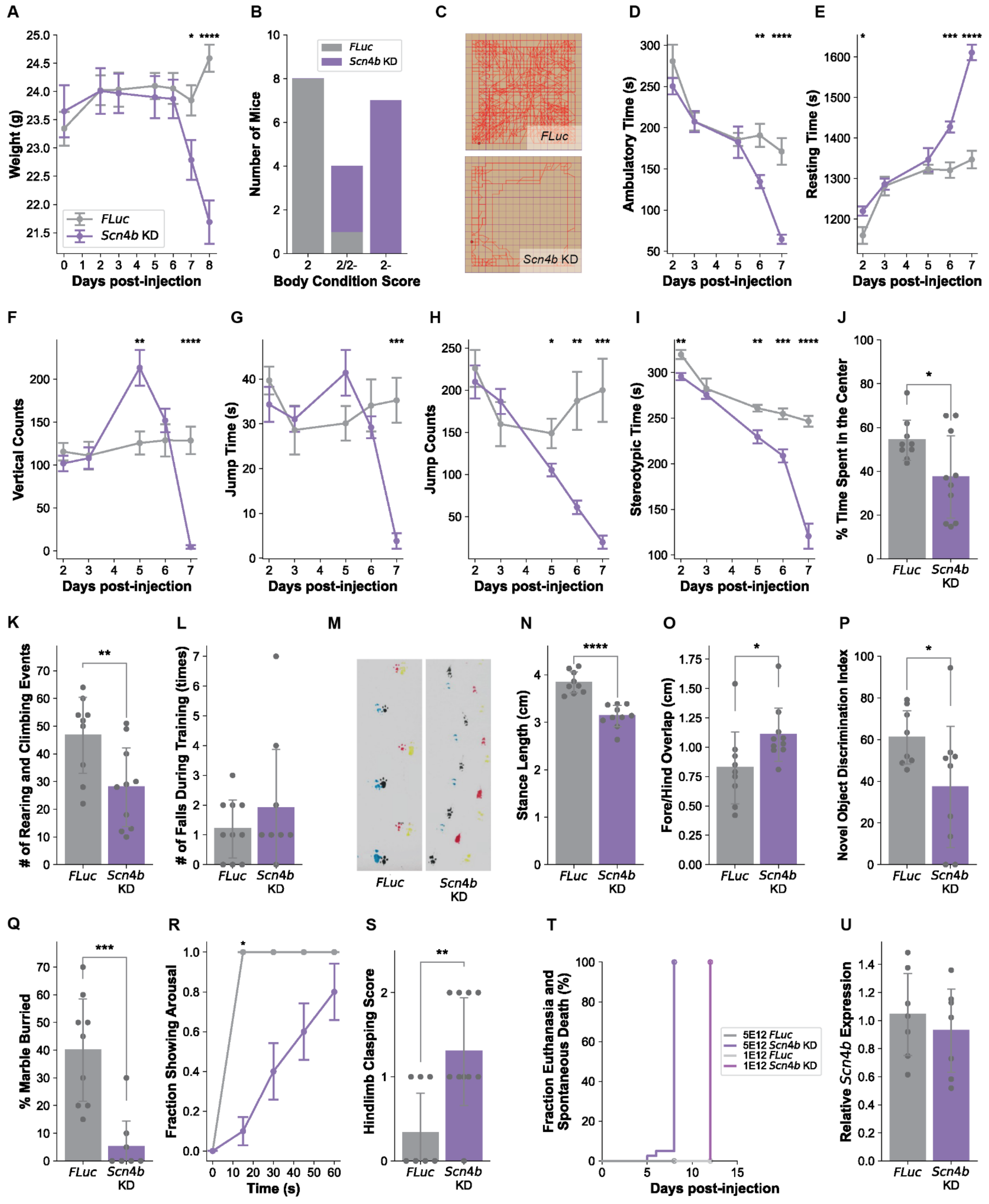

**Supplementary Figure 2. AAV PHP.eB hSyn CRISPR/CasRx Knockdown of *Scn4b* in adult wild-type mice promotes HD-related motor and coordination deficits. (A)** Body weight measurements comparing *Scn4b* KD mice and *FLuc* non-targeting control mice. **(B)** Body condition score. **(C)** Representative travel paths of a mouse in the open field test. **(D)** Ambulatory time, **(E)** resting time, **(F)** vertical counts, **(G)** jump time, **(H)** jump counts, **(I)** stereotypic time as measured by open field testing over a 30 min testing period. **(J)** Thigmotaxis analysis of horizontal time in open field testing on day 7. **(K)** Number of rearing and climbing events in the rearing and climbing test. **(L)** Number of falls during rotarod training. **(M)** Representative paw prints from gait analysis and the quantification of **(N)** stance and **(O)** overlap of the paw prints. **(P)** Novel object discrimination index. **(Q)** Percentage of marbles buried in the marble burying test. **(R)** Fraction of mice aroused following the opening of the home cage lid. **(S)** Hindlimb clasping score. Data are shown as mean ± SEM for all longitudinal time course data and mean ± SD for single-time-point comparisons (*n*=8-10 mice per group). One-sided t-test: *$p \leq 0.05$, **$p \leq 0.01$, ***$p \leq 0.001$, ****$p \leq 0.0001$. **(T)** Percent euthanasia and spontaneous death observed in our *in vivo* CasRx *Scn4b* KD. **(U)** Relative *Scn4b* RNA level in heart tissue, as determined by qPCR. Data are shown as mean ± SD. One-sided t-test: n.s.

### Supplemental Figure 3

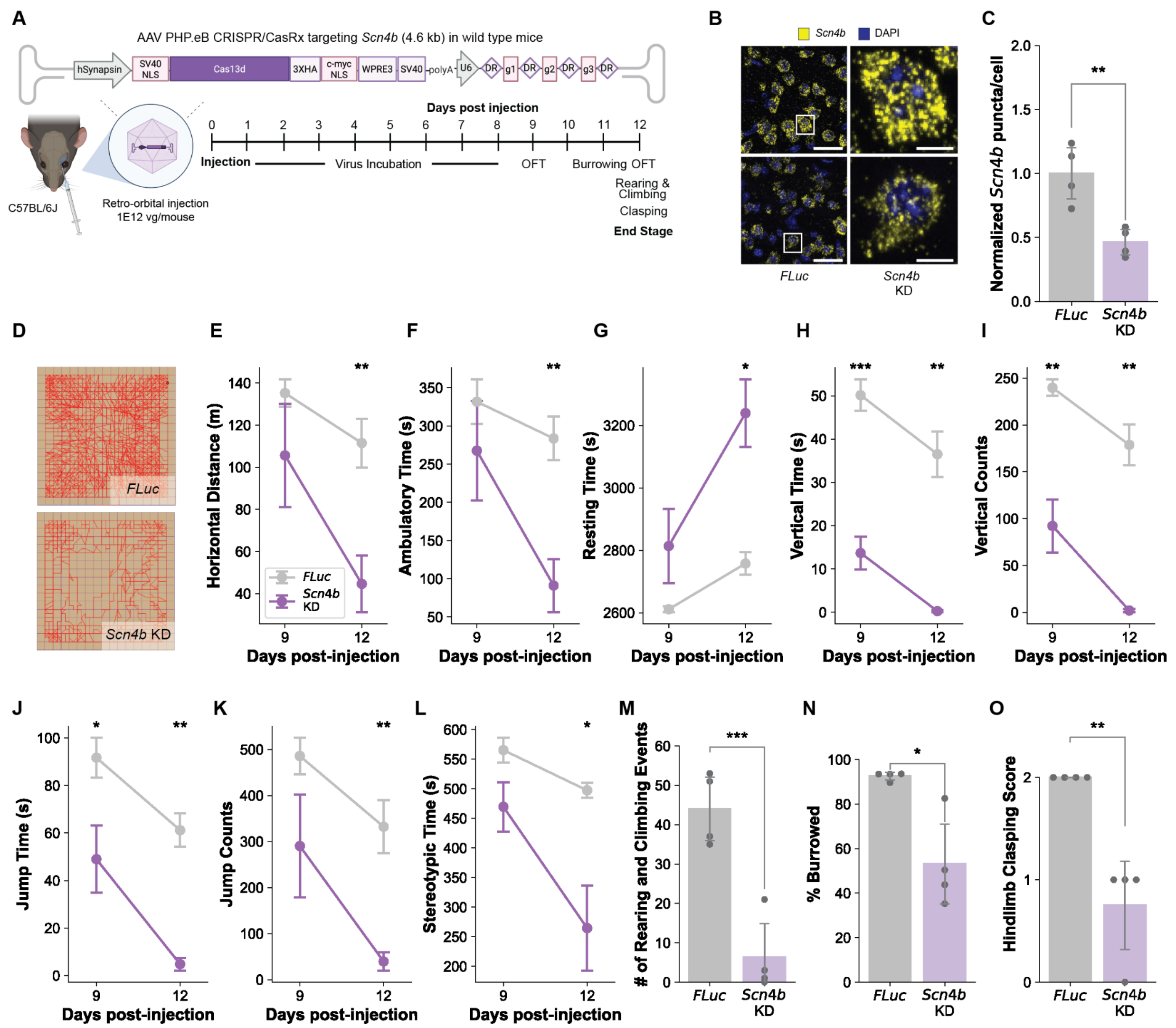


**Supplementary Figure 3. Low titer AAV PHP.eB hSyn CRISPR/CasRx Knockdown of *Scn4b* in adult wild-type mice promotes HD-related motor and coordination deficits. (A)** Schematic showing the *Scn4b* knockdown (KD) approach in adult wild-type mice and the timeline for behavioral testing. **(B)** Representative RNAScope *in situ* hybridization images of striatal tissue from mice injected with 1E12 vg/mouse of PHP.eB hSyn::CRISPR/CasRx containing guides targeting *Scn4b*, showing the reduction of *Scn4b* signal (yellow). *FLuc* denotes data from tissue with the guides targeting Firefly Luciferase, which is not present in the mouse genome. Scale bar, 25 μm (left) and 5 μm (right). **(C)** Relative quantification of *Scn4b* puncta count. Data are shown as mean ± SD (*n*=4 mice per condition; 25 images (63x) per mouse). One-sided t-test: $p = 0.0058$. **(D)** Representative travel paths of a mouse in the open field test. **(E)** Horizontal distance, **(F)** ambulatory time, **(G)** resting time, **(H)** vertical time and **(I)** count, **(J)** jump time and **(K)** count, **(L)** and stereotypic time traveled as measured by open field test over a 60 min testing period. **(M)** Number of rearing and climbing events in the rearing and climbing test. **(N)** Percentage of food pellets burrowed (by total mass) in the burrowing test. **(O)** Hindlimb clasping score. Data are shown as mean ± SEM for all longitudinal time course data and mean ± SD for single-time-point comparisons (*n*=4 mice per group). One-sided t-test: *$p \leq 0.05$, **$p \leq 0.01$, ***$p \leq 0.001$. Figure in panel A was created with BioRender.

### Supplemental Figure 4

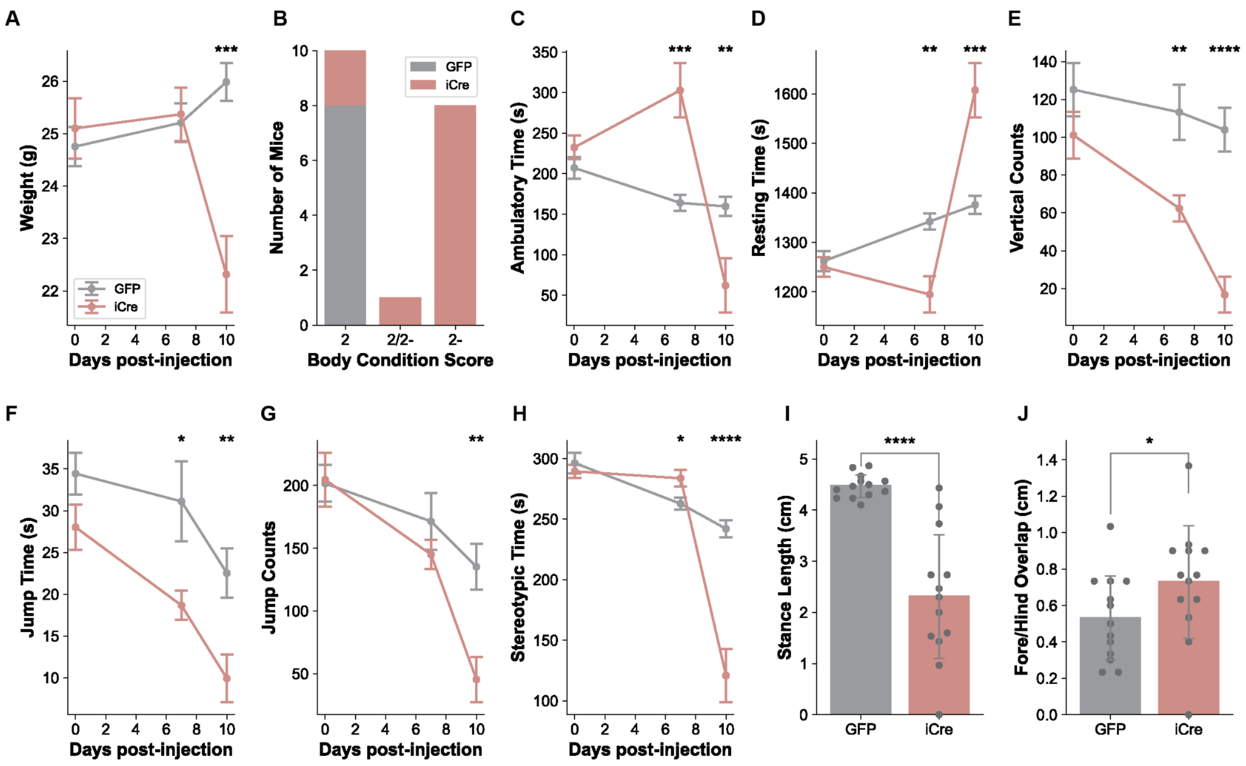


**Supplementary Figure 4. Conditional knockout of *Scn4b* in *Scn4b* floxed mice promotes HD-related motor and coordination deficits. (A)** Body weight measurements comparing *Scn4b* KO mice and GFP control. **(B)** Body condition score. **(C)** Ambulatory time, **(D)** resting time, **(E)** vertical time count, **(F)** jump time and **(G)** count, **(H)** and stereotypic time traveled as measured by open field test over a 60 min testing period. **(I-J)** Gait analysis quantification of stance and overlap of the paw prints. Data are shown as mean ± SEM for all longitudinal time course data and mean ± SD for single-time-point comparisons ($n$=13 mice per group). One-sided t-test: *$p \leq 0.05$, **$p \leq 0.01$, ***$p \leq 0.001$, ****$p \leq 0.0001$.

### Supplemental Figure 5

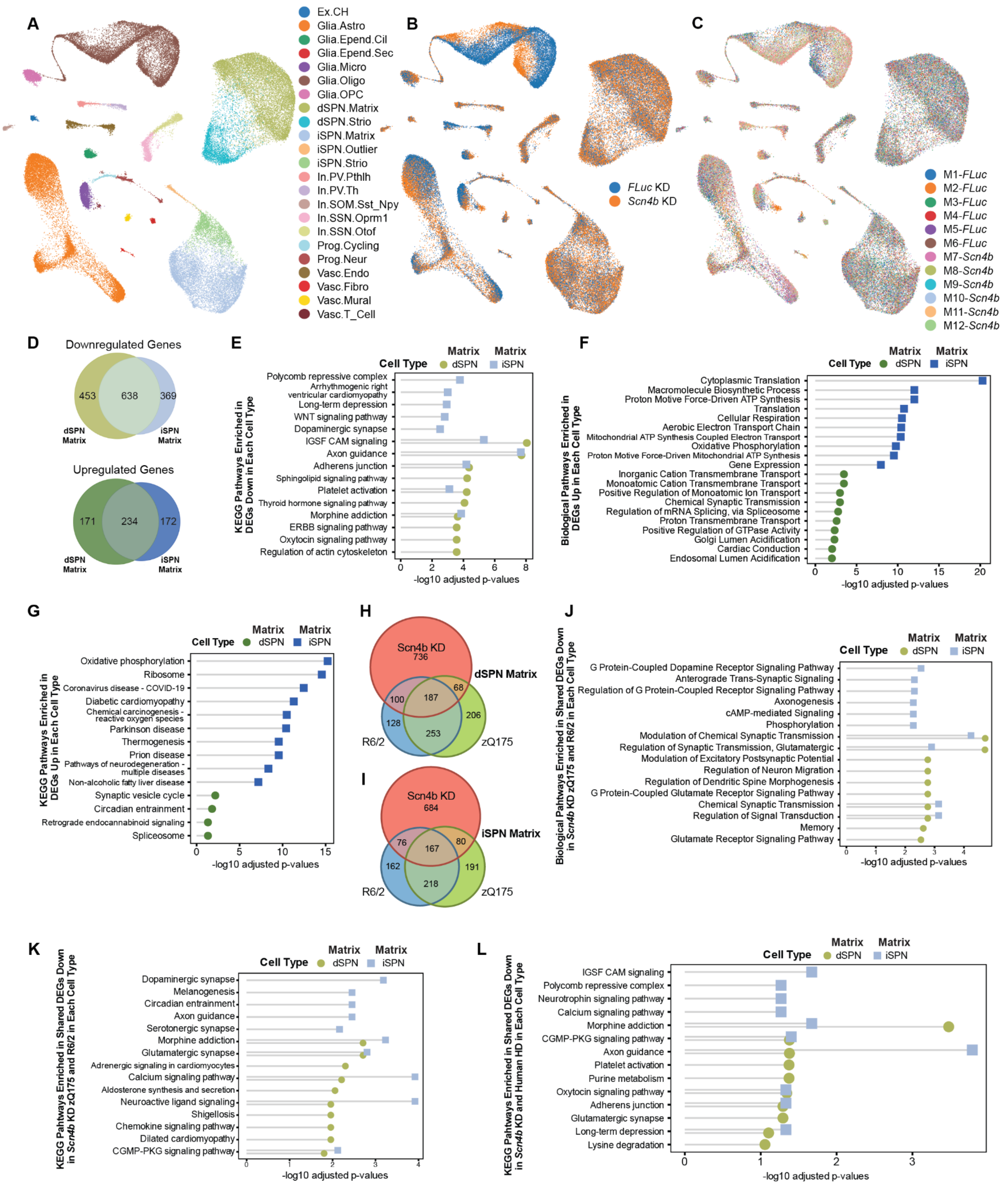

**Supplementary Figure 5. snRNA sequencing analysis of striatal cell types of *Scn4b* CRISPR/CasRx KD mice.**

**(A)** UMAP representing all cell types captured from striatal tissue of the *Scn4b* KD and control (*FLuc* KD) mice. UMAP showing the overlap between **(B)** *FLuc* and *Scn4b* KD knockdown conditions, and **(C)** individual mouse samples. **(D)** Venn diagram showing the overlap between matrix dSPNs and iSPNs upregulated and downregulated DEGs of *Scn4b* KD mice. **(E)** KEGG pathway analysis of downregulated DEGs in matrix dSPNs and iSPNs of *Scn4b* KD mice. **(F)** GOBP and **(G)** KEGG pathway analysis of upregulated DEGs in matrix dSPNs and iSPNs of *Scn4b* KD mice. Venn diagram showing the overlap between DEGs in matrix **(H)** dSPNs and **(I)** iSPNs between *Scn4b* KD and R6/2 and zQ175 HD mouse models.[1] **(J)** GO BP and **(K)** KEGG pathway enrichment analysis of downregulated DEGs in matrix dSPNs and iSPNs of the top overlapping modules among *Scn4b* KD and the R6/2 and zQ175 HD mouse models. **(L)** KEGG pathway enrichment analysis of the shared downregulated DEGs in matrix dSPNs and iSPNs from *Scn4b* KD mice and human HD.[2]

### Supplemental Figure 6

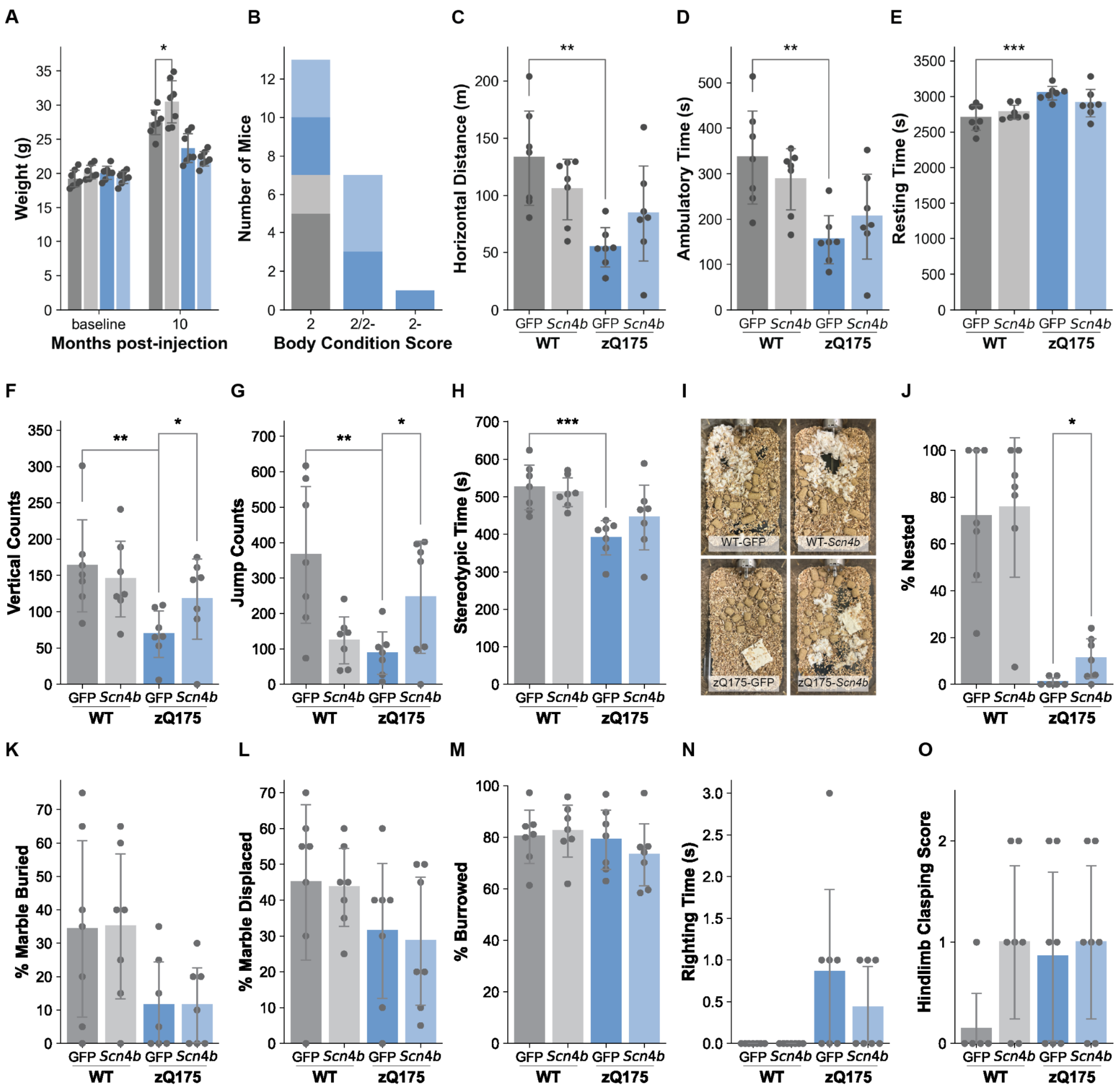


**Supplementary Figure 6. Striatal and motor-cortex-specific *Scn4b* overexpression in the zQ175 HD model mice rescues HD-related motor and coordination deficits.** **(A)** Body weight measurements. **(B)** Body condition score. **(C)** Horizontal distance, **(D)** ambulatory time, **(E)** resting time, **(F)** vertical counts, **(G)** jump counts, and **(H)** stereotypic time as measured by open field test over a 60 min testing period, for wild-type and zQ175 mice injected with 1E12 vg/mouse of either AAV PHP.eB *GPR88*::GFP or *GPR88*::*Scn4b* OX at 10 months post-injection (12 months of age). **(I)**

Representative nests after the overnight nesting test across experimental groups. **(J)** Percentage of nesting. Percentage of marbles **(K)** buried and **(L)** displaced in the marble burying test. **(M)** Percentage of food pellets burrowed (by total mass) in the burrowing test. **(N)** Righting reflex time. **(O)** Hindlimb clasping score. For all tests, data are shown as mean ± SD ($n$=7 mice per group). One-sided t-test: *$p \leq 0.05$.

### Supplemental Figure 7

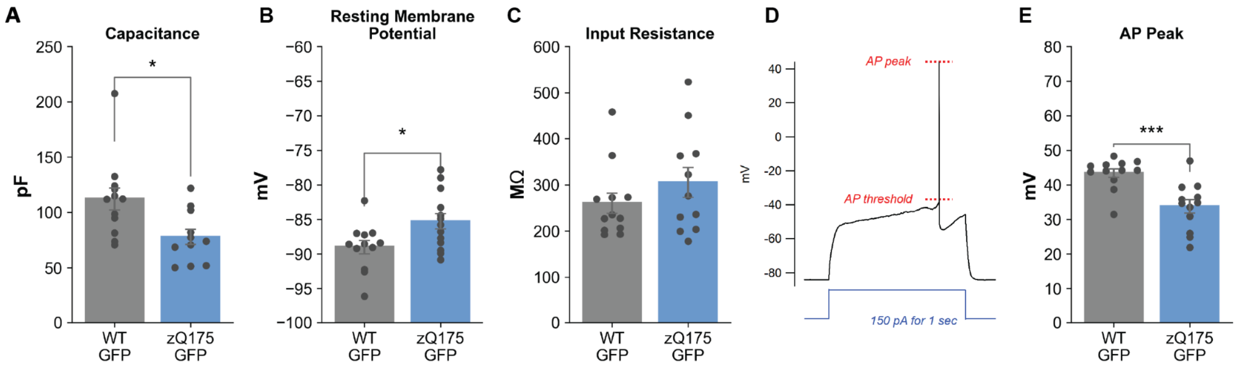


**Supplementary Figure 7. SPNs of wild-type and zQ175 mice show some differences in membrane properties.** **(A)** Capacitances, **(B)** resting membrane potentials, and **(C)** input resistances of WT-GFP OX compared to zQ175-GFP OX mice with individual values. Data are shown as mean ± SEM. One-sided t-test: *$p$ = 0.015 (capacitance) and 0.024 (resting membrane potential). **(D)** A representative trace of membrane potential showing an AP elicited by 1 sec-long current step of 150 pA. The two dotted lines (red) indicate the AP threshold and peak. **(E)** AP peaks for WT-GFP OX and zQ175-GFP OX mice. Data are shown as mean ± SEM. One-sided t-test: ***$p$ = 0.0006.

### Supplemental Figure 8

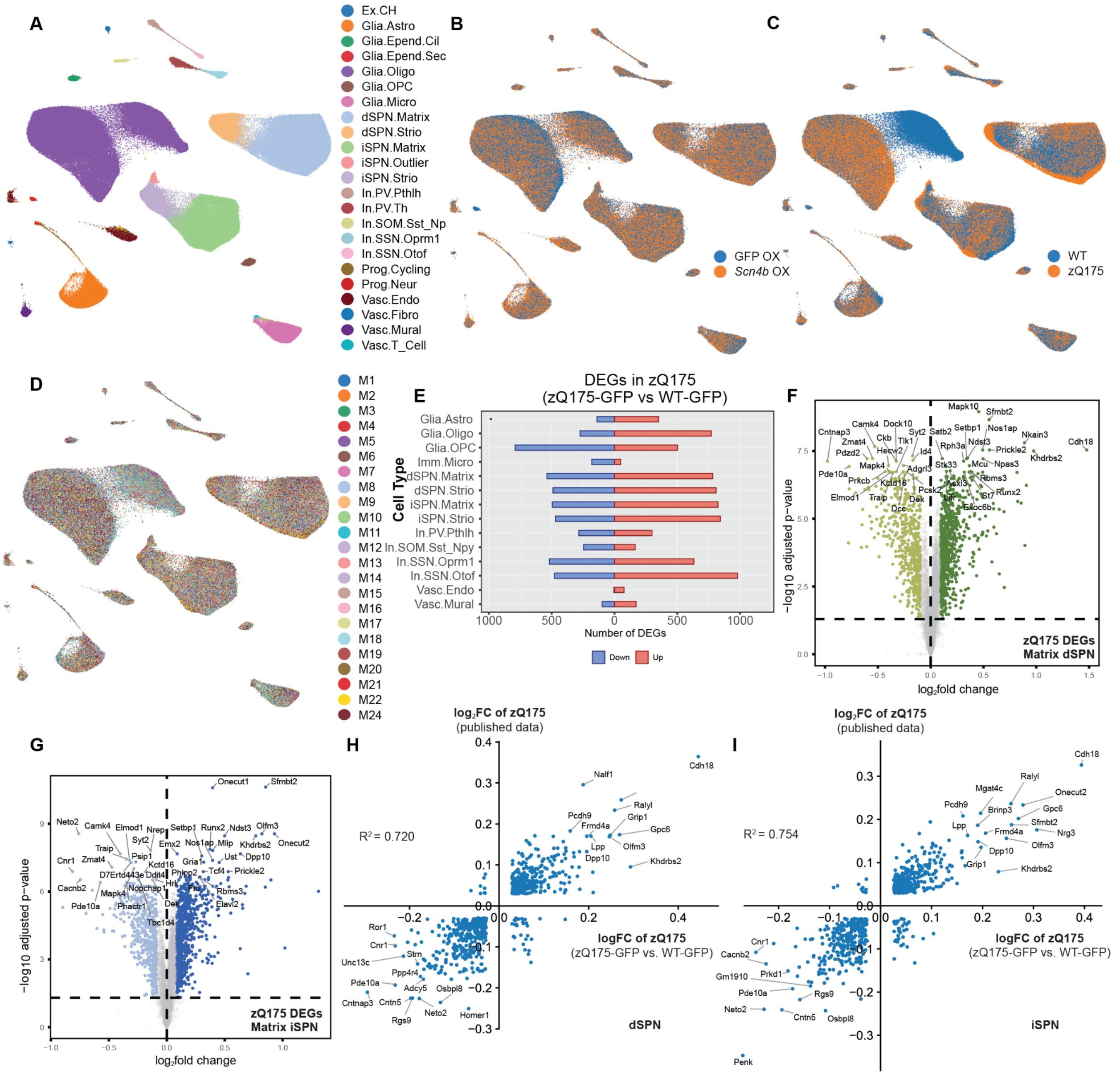


**Supplementary Figure 8. snRNA sequencing analysis of striatal cell types to reveal the zQ175 disease model effect at 13 months of age. (A)** UMAP representing all cell types captured from striatal tissue of all mice included in this OX study. UMAP showing the overlap between **(B)** GFP OX and *Scn4b* OX knockdown samples, **(C)** genotypes, and **(D)** individual mouse samples. **(E)** Back-to-back bar chart showing the number of genes that are dysregulated (DEGs) in the striatum of zQ175 mice (zQ175-GFP OX vs WT-GFP OX; $\log_2$ fold change abs($z$) >= 1 and $p$ adj. FDR < 0.05). Volcano plot showing these significant DEGs in matrix **(F)** dSPNs and **(G)** iSPNs of zQ175 mice. Quadrant plots showing $\log_2$ fold changes for the zQ175-GFP OX vs. WT-GFP OX comparison compared to published zQ175 data[1] for matrix (**H)** dSPNs and **(I)** iSPNs.

### Supplemental Figure 9

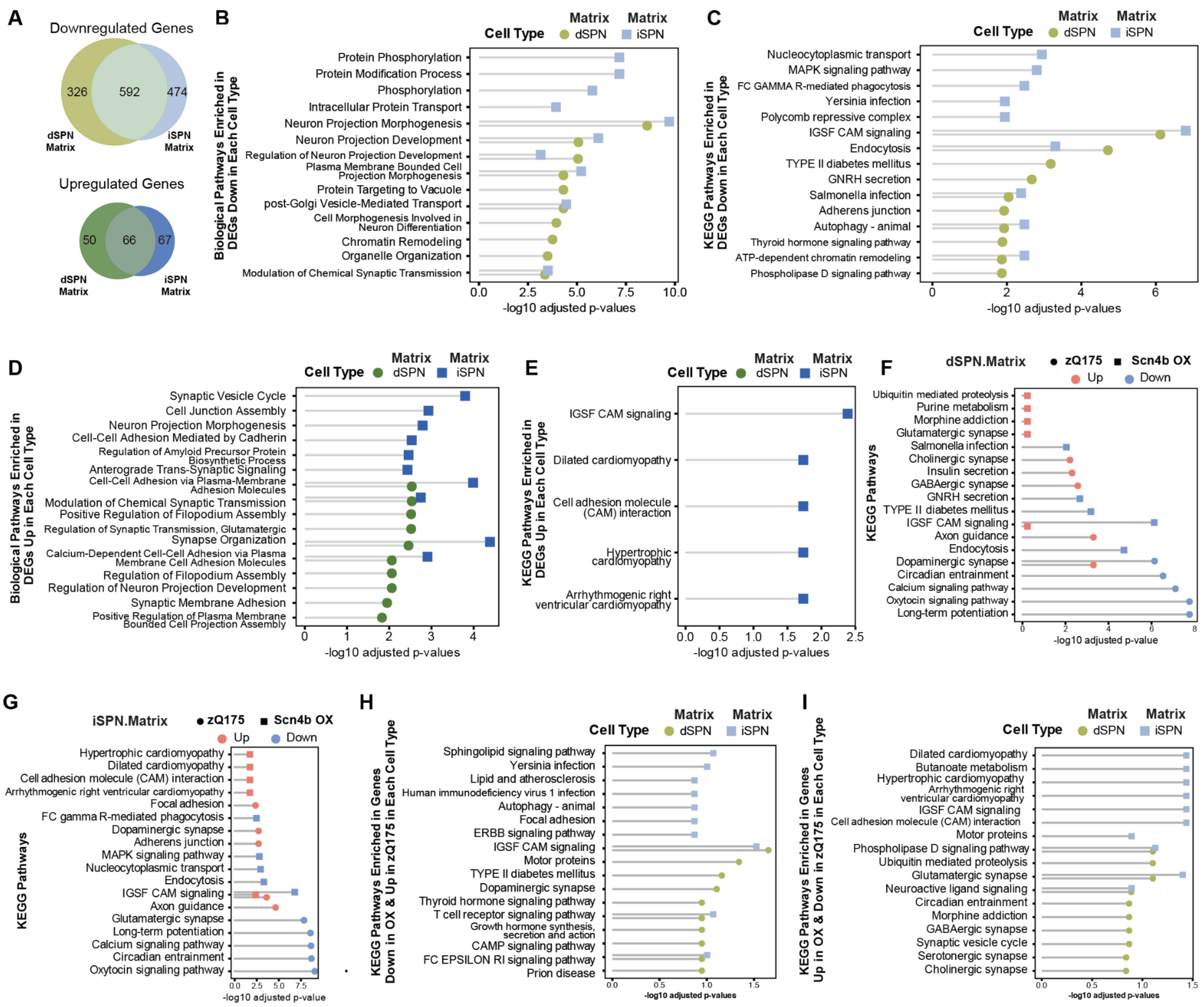


**Supplementary Figure 9. snRNA sequencing analysis of striatal cell types of zQ175 mice to determine the effect of *Scn4b* OX. (A)** Venn diagrams showing the overlap between matrix dSPNs and iSPNs, for upregulated and downregulated DEGs of the zQ175-*Scn4b* OX mice. **(B)** GO BP and **(C)** KEGG pathway enrichment analysis of *Scn4b* OX downregulated DEGs (zQ175-*Scn4b* OX vs. zQ175-GFP) in matrix dSPNs and iSPNs. **(D)** GO BP and **(E)** KEGG pathway enrichment analysis of *Scn4b* OX upregulated DEGs (zQ175-*Scn4b* OX vs. zQ175-GFP) in matrix dSPNs and iSPNs. KEGG pathway enrichment analysis showing the upregulated and downregulated pathways of matrix **(F)** dSPNs and **(G)** iSPNs across the zQ175 and *Scn4b* OX comparisons. KEGG pathway enrichment analysis of **(H)** genes that are downregulated in the *Scn4b* OX comparison and upregulated in the zQ175 comparison and **(I)** genes that are upregulated in the *Scn4b* OX comparison and downregulated in the zQ175 comparison.

### SUPPLEMENTAL REFERENCES


1. Matsushima, A., Pineda, S.S., Crittenden, J.R., Lee, H., Galani, K., Mantero, J., Tombaugh, G., Kellis, M., Heiman, M., and Graybiel, A.M. (2023). Transcriptional vulnerabilities of striatal neurons in human and rodent models of Huntington's disease. Nat. Commun. *14*, 282. https://doi.org/10.1038/s41467-022-35752-x.

2. Lee, H., Fenster, R.J., Pineda, S.S., Gibbs, W.S., Mohammadi, S., Davila-Velderrain, J., Garcia, F.J., Therrien, M., Novis, H.S., Gao, F., et al. (2020). Cell Type-Specific Transcriptomics Reveals that Mutant Huntingtin Leads to Mitochondrial RNA Release and Neuronal Innate Immune Activation. Neuron *107*, 891-908.e8. https://doi.org/10.1016/j.neuron.2020.06.021.

3. Obenauer, J.C., Chen, J., Andreeva, V., Aaronson, J.S., Lee, R., Caricasole, A., and Rosinski, J. (2022). Expression analysis of Huntington disease mouse models reveals robust striatum disease signatures. Preprint at Neuroscience, https://doi.org/10.1101/2022.02.04.479180.
